# Transcription factors read a second regulatory code in chromatin

**DOI:** 10.64898/2026.09.23.753737

**Authors:** Xin Zheng, Ying Tian, Jinyu Li, Haotian Sun, Xue Yue, Zhiyuan Xie, Litao Zheng, Yi Hui, Yuchen He, Danyang Zhou, Ying Xie, Xiaoqing Zhang, Lu Wang, Ke Xu, Yimeng Yin

## Abstract

Transcription factors (TFs) decode gene regulatory information written in DNA, yet how this vocabulary is interpreted within chromatin remains largely unexplored. Here, using an upgraded NCAP-SELEX platform, we systematically map the nucleosomal DNA recognition landscapes of 269 human TFs. We uncover a widespread, chromatin-dependent mode of sequence recognition: many TFs recognize motifs on nucleosomal DNA that are distinct from their canonical naked-DNA binding sites, revealing that nucleosome architecture encodes a second gene regulatory code in chromatin. Cryo-electron microscopy structures of TF-nucleosome complexes demonstrate that this second code is read through a combination of nucleosome-induced DNA deformation and direct protein-histone contacts. Functional analyses show that these chromatin-encoded motifs actively promote chromatin accessibility and drive cell-type-specific cis-regulatory activity *in vivo*. Strikingly, a single TF can deploy its canonical and nucleosome-derived motif repertoires to partition and govern entirely distinct physiological programs. Together, our findings establish that TFs interpret two complementary layers of genomic information: the primary DNA sequence and a second code written into the nucleosome architecture. This chromatin-encoded layer of regulatory information fundamentally expands our understanding of how TF specificity is achieved and how gene regulatory networks are wired in multicellular organisms.

## Main Text

Precise control of gene expression underlies cellular identity, development and adaptive responses to environmental cues. Central to this regulation are transcription factors (TFs), which interpret gene regulatory information encoded within *cis*-regulatory elements through recognition of specific DNA sequence motifs (*1, 2*). Extensive biochemical and genomic studies have established these sequence preferences as a fundamental regulatory code that guides TF occupancy and gene expression (*3–5*). However, this framework has largely been derived from studies of naked DNA, whereas genomic DNA in living cells exists within a highly organized chromatin environment.

Nucleosomes, the basic units of chromatin organization, profoundly influence TF binding by constraining DNA accessibility and imposing structural limitations on sequence recognition (*6–9*). Although most TFs preferentially bind nucleosome-depleted regions, pioneer factors possess the unique ability to engage nucleosomal DNA and initiate chromatin remodeling during development, lineage specification and cellular reprogramming (*10–14*). Structural studies have revealed that nucleosome organization can reshape DNA geometry, alter base accessibility and create new protein–histone interaction surfaces (*15–19*), raising the possibility that nucleosomes do more than restrict TF access—they may actively redefine the rules by which TFs interpret DNA sequences.

Consistent with this possibility, several pioneer TFs, including Oct4, Sox2, Klf4, GATA3 and FoxA proteins, recognize nucleosomal DNA through sequence preferences distinct from their canonical motifs identified on naked DNA (*10, 20–22*). These observations suggest that chromatin may encode additional gene regulatory information beyond the underlying DNA sequence. However, whether nucleosome-dependent recognition represents isolated examples or a general feature shared across the human TF repertoire remains unknown. Moreover, the molecular principles by which nucleosomes reshape TF specificity and the functional consequences of such alternative recognition modes remain poorly understood.

Here, we systematically map nucleosomal DNA recognition landscapes for 269 human TFs using an upgraded nucleosome consecutive-affinity-purification systematic evolution of ligands by exponential enrichment (NCAP–SELEX) platform. We identify widespread nucleosome-dependent binding specificities that differ from canonical DNA motifs, uncover distinct positional modes of TF recognition on nucleosomal DNA, and reveal structural principles underlying chromatin-dependent specificity through cryo-electron microscopy. Integrating genomic and functional analyses, we demonstrate that nucleosome-derived recognition sequences contribute to cell-type-specific regulatory activity and chromatin accessibility. Together, our findings establish that nucleosomes encode a second gene regulatory code in chromatin that is directly interpreted by TFs, expanding the framework by which gene regulatory information is encoded and decoded.

### A systematic atlas of TF recognition on nucleosomal DNA

Although the DNA-binding specificities of most human TFs have been extensively characterized on free DNA (*1, 23–25*), how nucleosome organization influences TF sequence recognition remains poorly understood. To systematically define TF recognition in the nucleosomal context, we optimized NCAP-SELEX (*26, 27*) by increasing library complexity, applying more stringent selection conditions and implementing an improved computational framework for motif discovery and refinement (**Fig. 1A**; see **Methods**).

**Fig. 1.**
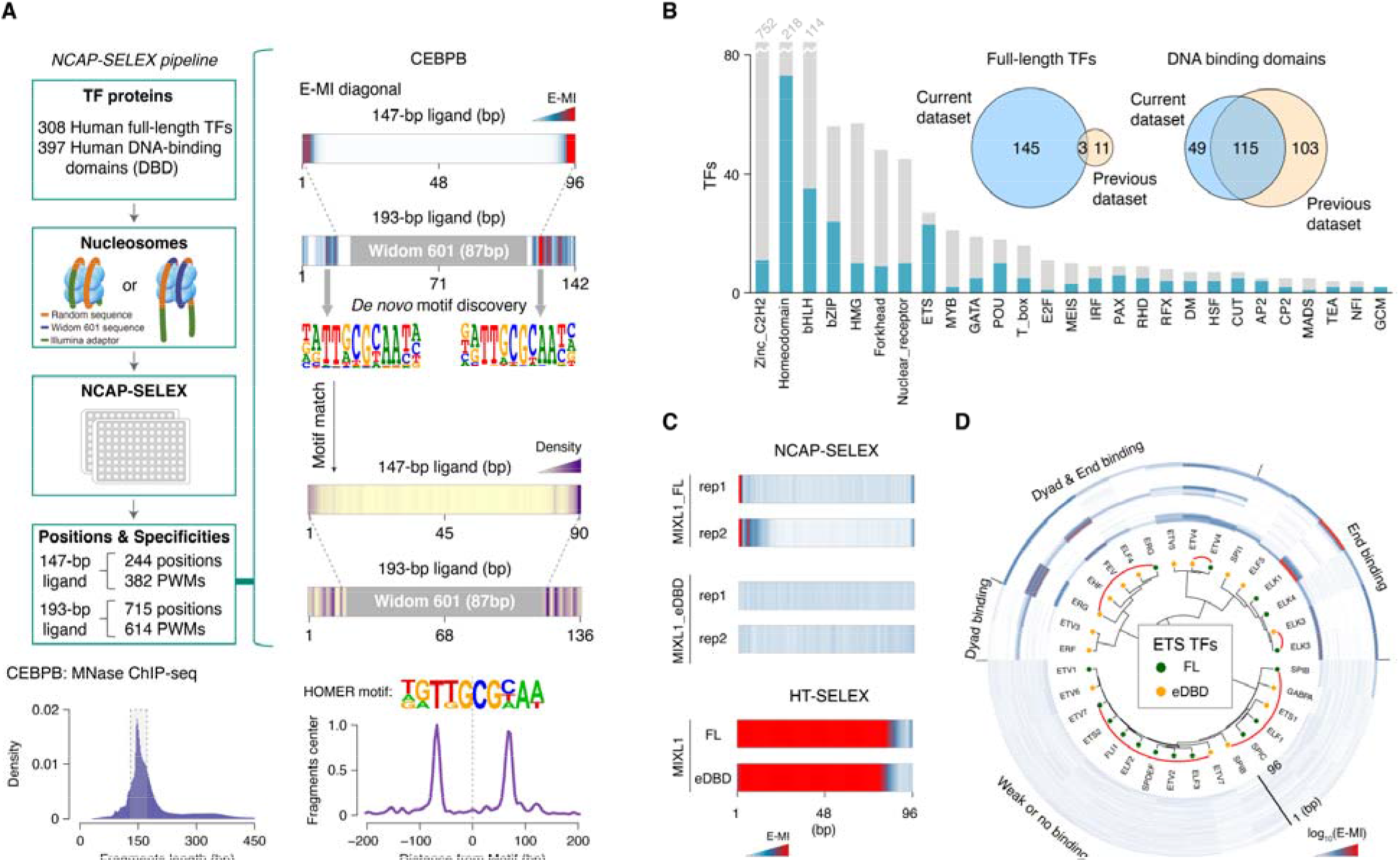
Comprehensive analysis of TF–nucleosome interactions by NCAP-SELEX. (**A**) NCAP-SELEX and *in vivo* validation. Top left: genome-scale NCAP-SELEX pipeline. Top right: example of TF-binding signal analysis by unsupervised E-MI (*26, 27*) and motif-based MOODS (*28*); representative NCAP-SELEX-derived PWM models are also shown. Bottom: *in vivo* validation of NCAP-SELEX results by MNase-ChIP-seq, showing fragment length distribution (left) and the enriched motif with its distance from fragment centers (right). (**B**) Coverage of TFs grouped by structural family. Blue bars represent TFs analyzed in this study; grey bars represent all known TFs (*1*). Inset: overlap percentage of full-length TFs (left) and DNA-binding domains (DBDs; right) between this study and our previous study (*26*). (**C**) E-MI diagonals of full-length MIXL1 and its extended DNA-binding domains (eDBDs) in NCAP-SELEX (top) and HT-SELEX (bottom) using the lig147. (**D**) Hierarchical clustering of E-MI diagonals of ETS factors in NCAP-SELEX using the lig147. E-MI diagonals for the same protein in its DBD (orange) and full-length (green) forms are connected by red arcs.

Using this platform, we profiled more than 700 active human TF proteins, including 308 full-length TFs and 397 extended DNA-binding domains (eDBDs) that displayed robust sequence-specific binding in high-throughput SELEX (HT-SELEX) assays (**table S1**). Together, these proteins represent approximately 40% of the high-confidence human TF repertoire (*1*). TFs were interrogated using two complementary nucleosome libraries: a canonical 147-bp nucleosome library (lig147), which permits systematic interrogation of binding across the nucleosomal DNA gyre, and a 193-bp library (lig193), in which an 87-bp Widom 601 positioning sequence flanked by randomized DNA enables analysis of TF binding near nucleosome entry–exit sites (**Fig. 1A** and **fig. S1**).

To identify TF binding motifs and their preferred positions on nucleosomal DNA, we integrated *de novo* motif discovery (Autoseed), enriched-sequence mutual information (E-MI) analysis and motif mapping using MOODS (*26, 28, 29*) (**Fig. 1A**). Across all screened proteins, reproducible sequence enrichment was detected for 269 distinct TFs, comprising 148 full-length proteins and 164 eDBDs (**Fig. 1B** and **table S2**). The median motif discovery success rate was 43%, representing substantially improved coverage over our previous systematic study (*26*) (**Fig. 1B**). Full-length TFs exhibited higher assay success rates than isolated DBDs (148/308 versus 164/397), suggesting that regions outside the DBD facilitate nucleosome recognition for a subset of TFs (*18, 19, 30*) (**Fig. 1C**).

Mapping TF occupancy across nucleosomal DNA reproduced the three predominant binding modes: binding near nucleosome entry–exit regions, binding around the dyad axis, and periodic binding to solvent-exposed DNA sites along the nucleosomal gyres (**Fig. 1A** and **fig. S1**). Together, these analyses establish a systematic map of human TF recognition on nucleosomal DNA, providing a foundation for investigating how nucleosome architecture shapes TF binding specificity.

### Nucleosome recognition diverges across TF families

Having established a global map, we next examined how nucleosome-binding properties vary across structural families. Entry–exit regions constituted the most permissive nucleosomal environment, with more than 40% of TFs binding lig193, indicating that most nucleosome-binding TFs preferentially recognize partially exposed DNA near nucleosome termini (**fig. S1** and **table S3**). In contrast, only 55 TFs accessed sequences near the center of lig147, demonstrating that the nucleosome interior represents a substantially more restrictive binding landscape.

Distinct TF structural families exhibited characteristic modes of nucleosome recognition. Members of the bHLH, bZIP, AP-2 and C2H2 zinc-finger families were largely confined to entry–exit regions, whereas RFX, NFAT and SOX family proteins efficiently recognized both entry–exit and dyad-associated sites (**fig. S1** and **table S3**). These observations indicate that different TF families possess markedly different capacities to engage nucleosomal DNA.

Notably, nucleosome recognition was not determined solely by DNA-binding domain conservation. Closely related paralogs frequently displayed distinct nucleosome-binding capacities and positional preferences (**Fig. 1D** and **fig. S1**). For example, within the ETS family, ELK1–4 and SPI1 (PU.1) predominantly recognized entry–exit regions, whereas ERG, FEV and ETV4 efficiently engaged both entry–exit and dyad-associated sites. By contrast, SPIC and SPDEF exhibited little or no detectable nucleosome-binding activity despite sharing highly conserved ETS domain (**Fig. 1D** and **fig. S1**). Similar divergence was observed in several additional TF families (**fig. S1**).

Together, these findings reveal that nucleosome recognition has diversified extensively both between and within TF families. Thus, the ability of TFs to engage nucleosomal DNA cannot be inferred solely from DNA-binding domain sequence or family classification, indicating that chromatin context introduces an additional layer of specificity beyond canonical DNA recognition.

### Nucleosomes reshape TF binding specificity

To determine whether nucleosomes alter TF sequence recognition, we compared motifs recovered from nucleosomal DNA with their corresponding motifs obtained from free DNA. Unexpectedly, whereas motifs enriched from free DNA closely matched previously reported monomeric or homodimeric binding models (*31*), a substantial fraction of nucleosome-derived motifs differed markedly from their canonical counterparts (**Fig. 2A**). From 996 position weight matrices (PWMs) recovered across 959 preferential binding positions for 312 TF proteins, a motif similarity network using Kullback–Leibler divergence (*23*) (KL) identified 268 previously unrecognized nucleosome-specific motifs, representing 142 distinct DNA-binding specificities (**fig. S2**, **table S2** and **S3**).

**Fig. 2.**
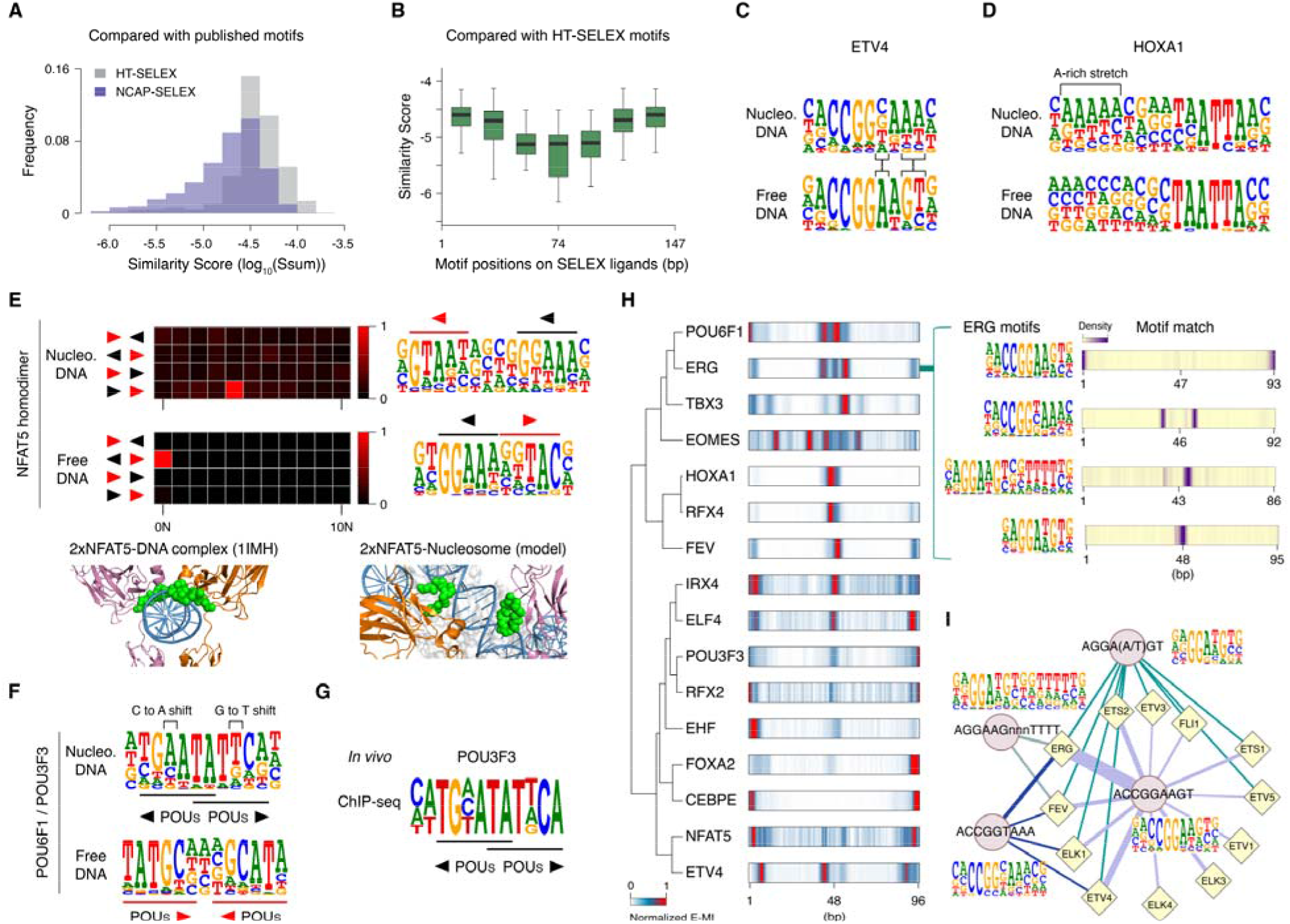
Characterization of nucleosome-embedded motifs identified by NCAP-SELEX. (**A**) Similarity of NCAP-SELEX (blue) and HT-SELEX (grey) motifs to JASPAR database motifs (*31*), measured by SSTAT (*57*). NCAP-SELEX motifs show substantially lower similarity to JASPAR motifs than HT-SELEX motifs do, indicating that a subset of NCAP-SELEX motifs are novel. (**B**) Similarity between HT-SELEX motifs and NCAP-SELEX motifs derived from different positions within the lig147 and lig193. (**C**) NCAP-SELEX (top) and HT-SELEX (bottom) motif logos for ETV4, representing a novel motif type. (**D**) NCAP-SELEX (top) and HT-SELEX (bottom) motif logos for HOXA1, representing a second novel motif type. (**E**) NCAP-SELEX (top) and HT-SELEX (bottom) motif logos for NFAT5. Heatmaps show the occurrence of NFAT5 5-mer subsequences TTTCC (black arrowhead) and ATTAC (red arrowhead) in the indicated orientations in NCAP-SELEX and HT-SELEX enriched libraries. The NFAT5–DNA complex structure (PDB entry 1IMH) shows DNA encircled by two NFAT5 proteins; a schematic model of the NFAT5–nucleosome complex indicates that two proteins target DNA with distinct spacing and orientation to avoid steric clashes with histone proteins. (**F**) NCAP-SELEX (top) and HT-SELEX (bottom) motif logos for POU6F1/POU3F3. The black arrowhead indicates the novel POU motif consensus (TATTC), and the red arrowhead indicates the canonical POU motif consensus (TATGC). (**G**) *In vivo* validation. The motif logo represents the POU3F3 motif, *de novo* identified by HOMER (*82*) from the 500 most enriched ChIP-seq peaks. (**H**) Hierarchical clustering of E-MI diagonals for TFs displaying diverse binding modes and specificities. Insets show four motif logos of ERG and motif-matching results on lig147, enriched at cycle 3 of NCAP-SELEX. Full data are shown in **fig. S4**. (**I**) Overview of class I ETS factor binding to distinct subsequences. Diamonds represent ETS proteins; circles represent enriched subsequences (*k*-mers). Edges connect proteins to k-mers that they enrich; edges are color-coded for clarity, and edge thickness represents enrichment strength. Motif logos show four distinct PWM models enriched by FEV.

These nucleosome-specific motifs exhibited distinct positional preferences. Unlike motifs that closely resembled canonical HT-SELEX motifs and were predominantly enriched near nucleosome entry–exit regions, nucleosome-specific motifs were concentrated within the nucleosome interior (SHL −3 to +3), around the dyad axis (**Fig. 2B**). Their preferential localization suggests that the structural environment of the nucleosome core—including DNA deformation and close proximity to the histone octamer—creates recognition interfaces that are inaccessible on free DNA.

To understand how TFs adapt sequence recognition to this chromatin environment, we compared each nucleosome-derived motif with its closest free-DNA counterpart, revealing four non-mutually exclusive modes of adaptation (**Fig. 2, C** to **F**, and **fig. S3**). Some TFs recognized truncated or degenerate versions of canonical motifs, thereby accommodating structural constraints imposed by the nucleosome (for example, POU2F2 and FIGLA; **fig. S3** and previous work (*10, 20–22*)). Others tolerated nucleotide substitutions within the core recognition sequence while maintaining sequence-specific contacts (for example, ETV4 and PAX7; **Fig. 2C** and **fig. S3**). A third group required AT-rich flanking sequences adjacent to otherwise canonical motifs (for example, HOXA1–2 and EOMES; **Fig. 2D**, and **fig. S3**). Finally, several TFs recognized dimeric motifs with altered spacing and orientation relative to their preferred free-DNA configuration, exemplified by NFAT5 (**Fig. 2E**). These recognition strategies frequently co-occurred within individual TFs.

POU-family factors illustrate this combinatorial adaptation. POU6F1 and POU3F3 recognized nucleosomal DNA near the dyad as homodimers through their POU_S_ domains, binding a tail-to-tail dimeric motif composed of overlapping TATTCA half-sites, whereas on free DNA they preferentially recognized a head-to-head TATGCA motif separated by one base pair (*25*) (**Fig. 2F**). Importantly, this nucleosome-specific motif was also detected in POU3F3 ChIP–seq datasets (**Fig. 2G**), supporting its physiological relevance. Moreover, several nucleosome-specific motifs were independently recovered in our previous NCAP-SELEX study (*26*) (**fig. S3**), demonstrating the reproducibility of these observations.

Together, these results demonstrate that nucleosomes do not simply restrict access to canonical TF binding sites but extensively remodel TF sequence recognition, generating a previously unrecognized repertoire of chromatin-specific binding motifs.

### Position-dependent sequence recognition by TFs on nucleosomes

Although most TFs recognize a population of sequences closely related to a single optimal site on free DNA, whether such sequence diversity expands when TFs engage nucleosomal DNA remains unclear. By systematically profiling individual TF binding specificities on nucleosomes, we found that dozens of TFs recognized two or more distinct sequences, resulting in multiple PWM models and diverse nucleosome-binding modes (**Fig. 2H**, **tables S2** and **S3**).

Strikingly, unlike TF binding on free DNA, where sequence recognition is largely determined by the intrinsic DNA-binding preference of each TF, nucleosomal binding specificity was strongly dictated by the precise position of the binding site within the nucleosome (**Fig. 2H** and **fig. S4**). Thus, the same TF can adopt distinct sequence preferences depending on its location relative to the nucleosome architecture.

This position-dependent adaptation is epitomized by the ETS family member ERG. While ERG recognizes only a single dominant motif on free DNA, it recognized four distinct motifs when bound to different positions within nucleosomal DNA (**Fig. 2H**). At the terminal entry–exit regions, ERG preferentially recognized its canonical ETS motif (5′-ACCGGAAGT-3′). In contrast, ERG binding at internal nucleosomal positions required three distinct, previously uncharacterized motifs—including ACCGGTAAA and AGGAAGNNNTTTT motifs enriched around the dyad axis (**Fig. 2H**). Moreover, these position-dependent binding modes were not uniformly shared across ETS family members. The canonical ETS motif was broadly recognized by class I ETS factors on nucleosomes, whereas the newly identified nucleosome-derived motifs displayed highly restricted TF preferences. For example, the ACCGGTAAA motif was recognized by a select subset of TFs including FEV, ELK1, ETV4, and ERG, while the AGGAAGNNNTTTT motif exhibited an even more stringent preference, engaged exclusively by FEV and ERG (**Fig. 2I**).

Together, these findings demonstrate that nucleosome architecture introduces a spatial layer of sequence specificity beyond intrinsic TF–DNA recognition, raising the fundamental question of how a single protein multi-tasks to read distinct codes across different nucleosomal positions.

### Molecular basis of position-dependent nucleosome recognition

To decipher how nucleosomal positioning reshapes TF sequence recognition at the atomic level, we determined cryo-electron microscopy (cryo-EM) structures of ERG bound to nucleosomes carrying the two representative novel motifs identified via NCAP-SELEX. The first structure contained the ACCGGTAAA motif positioned at SHL −1.7 (ERG-Nuc^ACCGGT^; 2.6 Å overall resolution), whereas the second contained the AGGAAG motif positioned at SHL −2.3 (ERG-Nuc^AGGAAG^; 3.2 Å overall resolution) (**Fig. 3, A** and **B**, **fig. S5** and **S6**, and **table S4**). These structures revealed that distinct nucleosomal positions impose unique geometric constraints on DNA, thereby driving alternative ERG recognition modes.

**Fig. 3.**
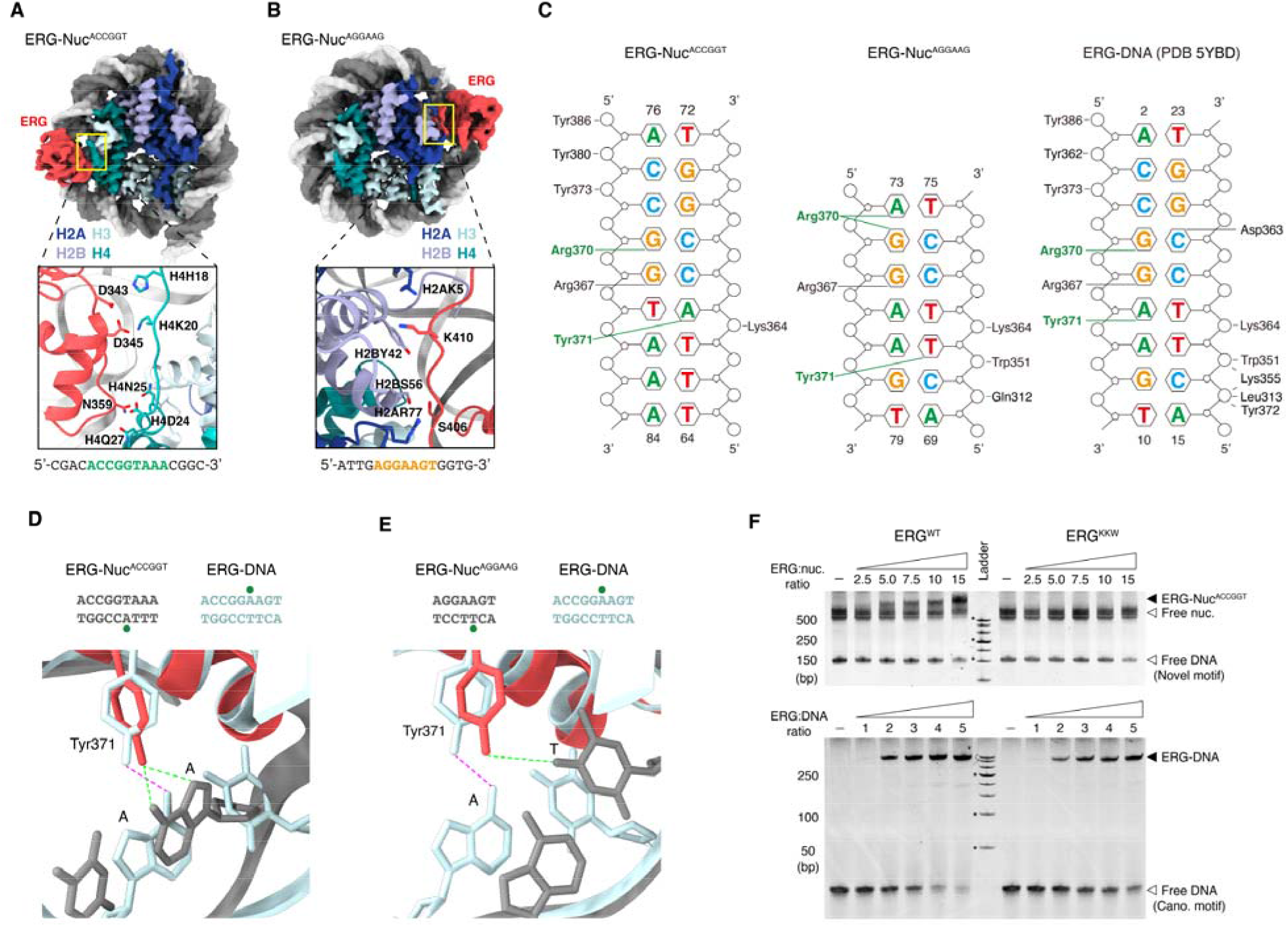
Mechanisms of novel motif recognition. (**A**) Cryo-EM structure of the ERG–Nuc^ACCGGT^ complex. Top: cryo-EM map of ERG–Nuc^ACCGGT^ at 2.6 Å resolution. Bottom: close-up view of contacts between ERG and the H4 tail in the structural model, with key residues shown as sticks. The DNA sequence recognized by ERG in the complex is shown below the structure and colored in green. (**B**) As in **A**, but for the ERG–Nuc^AGGAAG^ complex (3.2 Å resolution). The DNA sequence recognized by ERG in this complex is shown in orange. (**C**) Schematic overview of contacts formed by ERG with DNA in the ERG–Nuc^ACCGGT^ (left), ERG–Nuc^AGGAAG^ (middle), and ERG–DNA (right; PDB entry 5YBD) complexes. (**D**) Stereo view of ERG–DNA interactions mediated by Tyr^371^ with adenine bases in the ERG–Nuc^ACCGGT^ (ERG protein and DNA colored in red and grey, respectively) and ERG–DNA (ERG protein and DNA colored in cyan; PDB entry 5YBD) complexes. Hydrogen bonds are shown as dotted lines. The top panel shows the DNA sequences recognized by ERG in the ERG–Nuc^ACCGGT^ (left, grey) and ERG–DNA (right, cyan) complexes; dots indicate nucleotides that interact with Tyr^371^. (**E**) As in **D**, but for comparison between the ERG–Nuc^AGGAAG^ and ERG–DNA complexes. (**F**) Representative EMSA showing binding of increasing amounts of wild-type (ERG^WT^; left) and mutant (ERG^KKW^, D343K, D345K, and N359W; right) ERG to nucleosomes (top) and free DNA (bottom). Black arrowheads indicate ERG–nucleosome (top) and ERG–DNA (bottom) complexes; white arrowheads indicate free DNA and free nucleosomes. The DNA in the top panel contains the novel motif consensus (ACCGGTAAA), whereas the DNA at the bottom panel contains the canonical motif consensus (ACCGGAAGT).

In the ERG-Nuc^ACCGGT^ structure, the ETS domain adopted a global orientation similar to that observed in the ERG–DNA complex (*32*), inserting its recognition helix (α3) into the solvent-exposed major groove (**Fig. 3A**, **fig. S5** and **S7**). However, nucleosomal DNA itself exhibited pronounced structural deformation, including major-groove narrowing and an approximately 70° DNA bend away from the ERG interface. These DNA distortions altered the base contacts without requiring substantial protein conformational rearrangement. Specifically, Tyr^371^, which recognizes adenine within the canonical ETS core (5′-GGAA-3′) on free DNA, was repositioned to contact the corresponding base on the opposite DNA strand, resulting in recognition of a distinct core sequence (5′-GGTA-3′) (**Fig. 3, C** and **D**, and **fig. S7**).

Nucleosome incorporation also altered the energetic basis of ERG binding. Compared with the free DNA complex, ERG lost several phosphate-backbone interactions that normally contribute substantially to ETS–DNA affinity (*33*) (**Fig. 3C** and **fig. S7**). This reduction was compensated by additional interactions between the ERG α2 helix and the N-terminal tail of histone H4, which adopts a trajectory toward DNA near SHL −2 (**Fig. 3A** and **fig. S7**). Consistent with this structural model, mutation of ERG α2 residues involved in H4 interaction (Asp^343^, Asp^345^ and Asn^359^; ERG^KKW^) selectively impaired nucleosome binding while minimally affecting binding to free DNA with the canonical ETS motif (**Fig. 3F**). These results demonstrate that nucleosome-specific binding modes arise from a combination of altered DNA readout and histone-mediated stabilization.

We next examined ERG binding at SHL −2.3, where the AGGAAG motif was preferentially recognized. In this configuration, the ETS domain underwent an approximately 120° rotational repositioning relative to the ERG-Nuc^ACCGGT^ structure, allowing the recognition helix to engage a distinct solvent-exposed major groove (**Fig. 3B**, **fig. S6** and **S7**). Binding at this position was accompanied by local DNA unwrapping, with disruption of histone–DNA contacts between SHL +4.5 and SHL +7 (**fig. S7**), indicating that ERG exploits dynamic nucleosome features to access alternative recognition sites.

At this nucleosomal position, ERG retained a largely unchanged protein conformation compared with the free DNA complex (RMSD = 0.9 Å), whereas the surrounding DNA architecture was substantially remodeled (**fig. S7**). The altered DNA geometry redirected interactions involving Tyr^371^ and Arg^370^: Tyr^371^ was positioned adjacent to the thymine opposite the second adenine of the canonical 5′-GGAA-3′ core, whereas Arg^370^, which normally contacts the first guanine in free DNA, was repositioned to coordinate both the core guanine and an upstream adenine (**Fig. 3, C** and **E**, and **fig. S7**). Thus, similar to ERG-Nuc^ACCGGT^, alternative motif recognition at this position is primarily driven by nucleosome-induced DNA deformation rather than by major TF rearrangement.

Additional interactions further stabilized ERG binding in the nucleosomal context. These included phosphate-backbone contacts with flanking DNA regions, contacts between the ERG intrinsically disordered region (IDR; residues 399–414) and histones H2A/H2B, and proximity of the H2B N-terminal tail to ERG (**Fig. 3B** and **fig. S7**). However, IDR deletion had little effect on ERG–nucleosome binding in electrophoretic mobility shift assay (EMSA) assays, suggesting that these contacts are only partially contributory and that ERG recognition is mediated by a distributed network of DNA and histone interactions (**fig. S7**).

Together, these structures reveal a general mechanism by which nucleosomal architecture expands TF binding specificity. Rather than simply restricting TF access, nucleosomes reshape the DNA recognition landscape by altering DNA geometry, exposing distinct sequence contexts, and providing histone interaction surfaces. Consequently, a single TF can recognize multiple position-dependent motifs through alternative combinations of DNA readout and nucleosome-mediated stabilization.

### Nucleosome-specific motifs preferentially function during chromatin opening

To investigate the potential regulatory functions of nucleosome-derived novel motifs, we first examined their association with human *cis*-regulatory elements using candidate *cis*-regulatory elements (cCREs) from the cis-regulatory atlas (CATlas) (*34*). Novel motif matches were frequently enriched in cell-type-specific cCREs, with 203 of 268 novel motifs showing significant enrichment in one or a limited number of cell-type-specific cCRE sets (FDR < 0.01; see **Methods**), whereas the remaining motifs exhibited broader enrichment patterns across multiple cell types. This distribution was comparable to that observed for the corresponding canonical motifs (155/179; **Fig. 4A**, **fig. S8** and **table S5**), suggesting that nucleosome-derived motifs retain regulatory specificity despite their distinct sequence preferences. Moreover, the cell-type enrichment profiles of novel motifs closely resembled those of their canonical counterparts and were consistent with the known biological functions of the corresponding TFs (**Fig. 4B** and **fig. S8**). For instance, both novel and canonical ERG motifs were preferentially enriched in endothelial and lymphocyte cCREs, consistent with the established roles of ERG in endothelial homeostasis and definitive hematopoiesis (*35, 36*). Similarly, POU6F1-associated novel and canonical motifs showed strong enrichment in neuronal and microglial cCREs, consistent with the function of POU6F1 in neuronal development and plasticity (*37*).

**Fig. 4.**
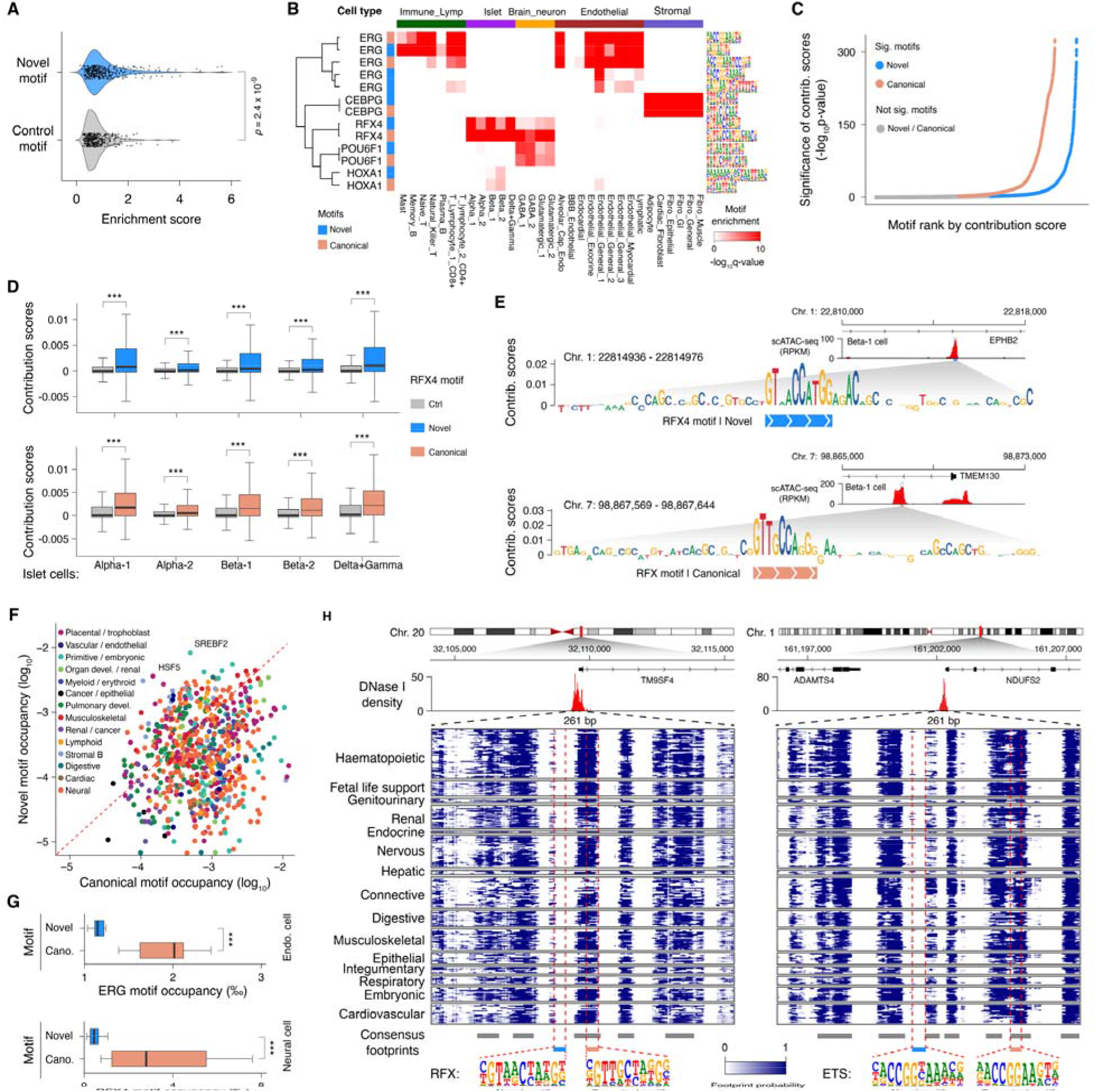
*In vivo* functional characterization of the novel motifs. (**A**) Enrichment of novel (blue) and control (grey) motifs in cell-type-specific cCREs from CATlas (*34*). For each motif, the top-three enrichment scores (−log_10_(q values)) across distinct cell types are shown. Statistical significance was determined by a two-sided Student’s *t*-test. (**B**) Hierarchically clustered heatmap showing enrichment of representative novel (blue) and canonical (orange) motifs in cell-type-specific cCREs from CATlas (*34*). Rows represent motifs; columns represent cell types. Full data are shown in **fig. S8**. (**C**) Distribution of contribution scores of novel and canonical motifs to chromatin accessibility. Motifs that significantly contribute to accessibility compared with their respective control motifs (adjusted P < 0.01, one-sided Wilcoxon rank-sum test) are colored in blue (novel) and orange (canonical). Motifs without significant contribution in any analyzed cell type (*34*) are shown in grey. (**D**) Example showing that novel and canonical motifs of a TF significantly contribute to chromatin accessibility across islet cell types. Center lines indicate medians; boxes indicate interquartile ranges (IQR); whiskers indicate 5th and 95^th^ percentiles of contribution scores for nucleotides matching novel (blue), canonical (orange), or control (grey) motifs within the top 30,000 ATAC-seq peaks. Statistical significance was determined by a one-sided Wilcoxon rank-sum test; ***P < 0.001. (**E**) Examples of the EPHB2 (top) and TMEM130 (bottom) loci in Beta-1 cells, showing observed chromatin accessibility, predicted per-base contribution scores, and annotated motif instances. (**F**) TF occupancy at canonical motif matches (x-axis) versus novel motif matches (y-axis) within DNase I hypersensitive sites (DHSs) across 243 ENCODE biosamples (*44*). (**G**) Examples of TF occupancy at novel (blue) and canonical (orange) motif matches within DHSs of endothelial (top) and neural (bottom) cells. Center lines indicate medians; boxes indicate IQR; whiskers indicate most extreme values within 1.5 × IQR. Statistical significance was determined by a two-sided Wilcoxon rank-sum test without multiple-testing correction; ***P < 0.001. **(H**) Heatmap of footprint posterior probabilities within two DHSs. Rows represent 243 ENCODE biosamples (*44*) grouped by tissue or organ systems; columns represent individual nucleotides. Grey boxes indicate consensus footprints in one or more cell or tissue types (footprint posterior > 0.99); blue and orange boxes indicate novel and canonical motif matches, respectively. The top panel shows windowed DNase I cleavage density, and the bottom panel shows representative motif logos for RFX4 (RFX family) and ERG (ETS family).

To determine whether novel motif sequences actively contribute to the establishment of accessible chromatin states, we next leveraged deep learning models trained to predict chromatin accessibility from DNA sequence (*38, 39*). We trained five-fold cross-validated ChromBPNet models for 110 cell types in the CATlas dataset (550 models in total) to predict both total ATAC-seq signal intensity and base-resolution read distributions within 1-kb regions centered on pseudobulk ATAC-seq peaks (**table S6**; see **Methods**). The trained models achieved a median Pearson correlation coefficient of 0.66 between predicted and observed read counts in held-out ATAC-seq peaks (**fig. S8** and **table S6**), demonstrating robust predictive performance.

We then applied DeepLIFT (*40*) to these models to quantify the contribution of individual nucleotides within cCREs to predicted accessibility profiles. This analysis revealed that sequences matching a substantial fraction of novel motifs (84/268, FDR < 0.01) exhibited significantly higher contribution scores than matched control sequences in cCREs from specific cell types or related cellular lineages (**Fig. 4C** and **table S6**; see **Methods**). Similar accessibility-promoting effects were observed for canonical motifs (95/179, FDR < 0.01; **Fig. 4C**), indicating that both motif classes encode sequence information relevant to chromatin regulation. For example, both novel and canonical RFX4 motif matches showed strong predicted contributions to chromatin accessibility in pancreatic islet cell cCREs, consistent with the established role of RFX family members as key regulators of islet cell development (*41, 42*) (**Fig. 4, D** and **E**).

Having established that novel motifs are associated with regulatory elements and contribute to predicted chromatin accessibility, we next asked whether these motifs are directly recognized by TFs within active *cis*-regulatory elements. To address this question, we analyzed ENCODE DNase I cleavage profiles, which provide nucleotide-resolution maps of TF occupancy in native chromatin contexts (*43, 44*). Compared with canonical motif matches, novel motif matches generally exhibited reduced TF occupancy within DNase I hypersensitive sites (DHSs) across 243 human cell and tissue types and states (*44*) (**Fig. 4F** and **table S7**). This trend was observed for most TF families, including RFX and ETS factors (**Fig. 4, G** and **H**), suggesting that although nucleosome-derived novel motifs can contribute to regulatory element activity, they are less frequently used as direct TF-binding sites after chromatin opening.

Nevertheless, several notable exceptions were identified. A subset of novel motifs, including those associated with HSF5 and SREBF2, displayed higher TF occupancy than their corresponding canonical motifs (**Fig. 4F** and **table S7**). These cases may reflect reduced sensitivity to CpG methylation due to altered sequence composition, as CpG methylation can strongly influence TF binding, or may represent previously unrecognized monomeric binding modes for TFs traditionally characterized as homodimeric factors (**fig. S3**).

Together, these findings indicate that nucleosome-derived novel motifs represent functional regulatory sequences that are preferentially incorporated into cell-type-specific *cis*-regulatory landscapes and contribute to the establishment of accessible chromatin states. However, unlike canonical motifs that frequently mediate direct TF recruitment on accessible DNA, most nucleosome-derived novel motifs appear to function primarily during chromatin opening and are subsequently dispensable for later transcriptional processes, such as preinitiation complex assembly.

### Biological roles of TF–novel-motif recognition *in vivo*

To determine whether recognition of nucleosome-derived novel motifs directs distinct regulatory functions *in vivo*, we performed structure-guided mutagenesis based on our cryo-EM models. We utilized the ERG^KKW^ mutant, which selectively disrupts key interactions with the histone H4 tail at SHL −1.7, thereby impairing nucleosomal novel motif recognition while preserving canonical ETS motif binding on free DNA (**Fig. 3F**). We then performed ATAC–seq in human embryonic stem cells (ESCs) expressing either wild-type ERG (ERG^WT^) or the ERG^KKW^. Expression of ERG^WT^ induced robust chromatin accessibility at genomic regions enriched for both canonical and novel ERG motifs, whereas ERG^KKW^ failed to open a subset of these regulatory elements (**Fig. 5, A** and **B**, and **fig. S9**).

**Fig. 5.**
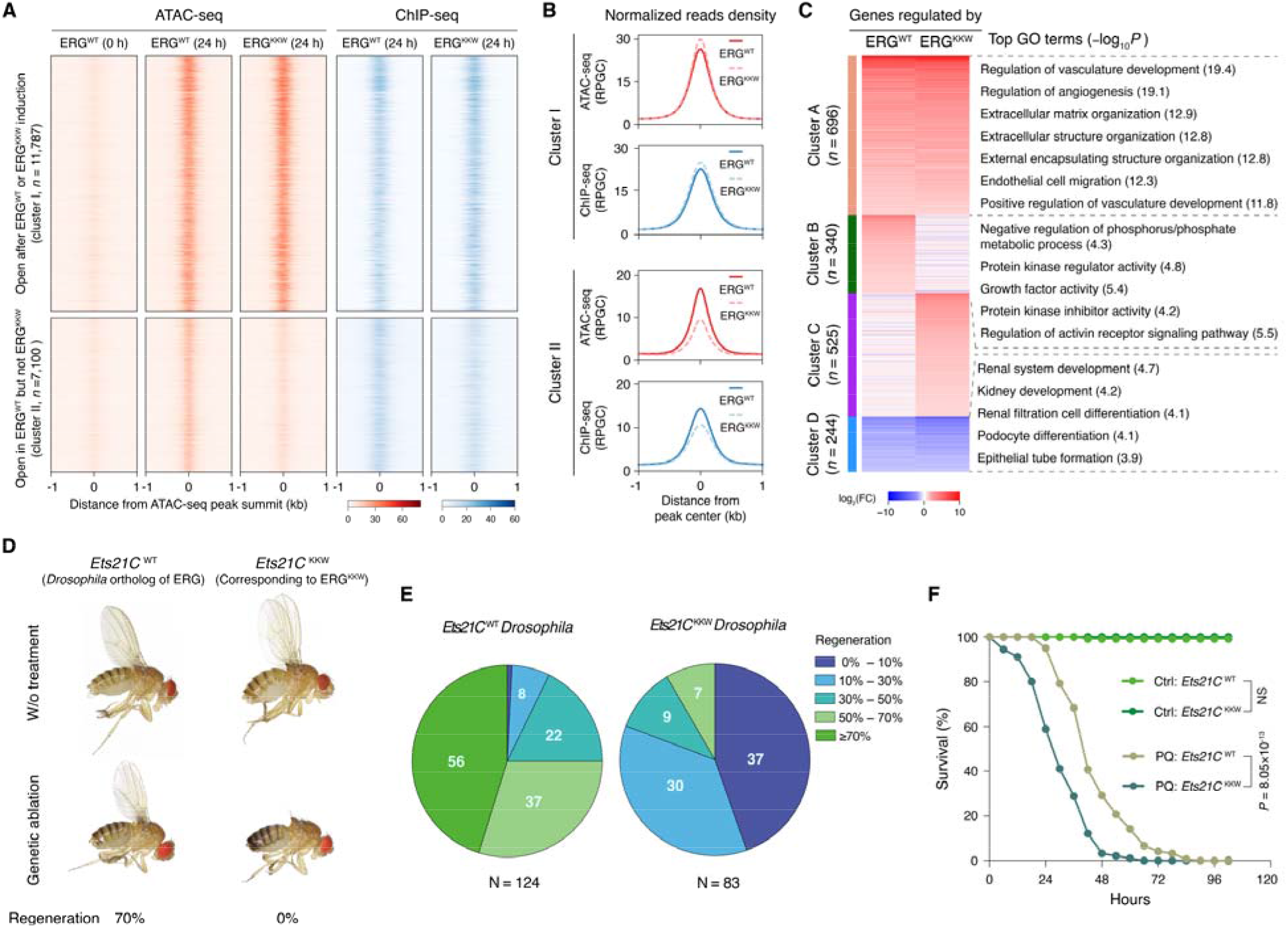
Biological roles of TF-novel-motif recognition. (**A**) Read-density heatmaps showing normalized ATAC-seq signal (red) and ChIP-seq intensity (blue) for ERG^WT^ and ERG^KKW^ in ESCs. ATAC-seq signals before and 24 h after ectopic induction of ERG^WT^ and ERG^KKW^ are shown. Chromatin sites are ranked by ATAC-seq signal in ERG^WT^-expressing ESCs and divided into two clusters: cluster I, sites that become open after ERG^WT^ or ERG^KKW^ expression; and cluster II, sites that become open only after ERG^WT^ expression. Cluster sizes and color scales are indicated. (**B**) Line plots showing average ATAC-seq (red) and ChIP-seq (blue) signals at clusters I and II. Solid and dashed lines indicate signals from ESCs expressing ERG^WT^ and ERG^KKW^, respectively. RPGC (reads per genomic content; 1× normalization). (**C**) k-means clustering (*k* = 4) of genes regulated by ERG^WT^ and/or ERG^KKW^. The top enriched biological process GO terms identified are shown adjacent to the corresponding clusters. (**D**) Extent of wing imaginal disc regeneration following genetic ablation in *Drosophila* expressing wild-type Ets21C^WT^ (the *Drosophila* ortholog of ERG) or mutant Ets21C^KKW^ (D287K, D289K, and N303W, corresponding to ERG^KKW^). Top panels show phenotypes of *Ets21C*^WT^ and *Ets21C*^KKW^ flies maintained under standard conditions; bottom panels show representative regeneration phenotypes for *Ets21C*^WT^ (≥70% regeneration) and *Ets21C*^KKW^ (no regeneration capacity) flies after genetic ablation assays. (**E**) Distribution of wing imaginal disc regeneration scores in *Ets21C*^WT^ (left) and *Ets21C*^KKW^ (right) *Drosophila*. Regeneration was scored by binning wings into five categories (0% − 10%, 10% − 30%, 30% − 50%, 50% − 70%, or ≥70%) and was calculated per population. Experiments were performed on separate days with a minimum of two vials per genotype and four replicates per genotype. (**F**) Survival of adult *Ets21C*^WT^ and *Ets21C*^KKW^ *Drosophila* fed with 50 mM paraquat (PQ) or mock solution (Ctrl). Survival curves were derived from four independent experiments. Statistical significance was determined by the log-rank test. NS, not significant.

To capture the temporal dynamics of this process, we mapped ERG occupancy by ChIP–seq 24 h after induction. ERG preferentially engaged genomic regions that were initially inaccessible, with over 60% of bound sites located within closed chromatin enriched for both canonical and novel motifs (**Fig. 5, A** and **B**, and **fig. S9**). This rapid engagement followed by gradual chromatin opening directly mirrors classical pioneer transcription factor activity (*12*), where DNA binding precedes chromatin remodeling. Notably, a substantial fraction of ERG-bound sites remained inaccessible 48 h after induction, further supporting the gradual nature of chromatin opening following pioneer factor recruitment (**fig. S9**).

We next investigated whether nucleosome-derived motif recognition regulates gene programs distinct from those controlled by canonical ERG motif. RNA–seq and gene ontology analyses revealed that ERG^WT^- and ERG^KKW^-responsive genes both governed vascular development, angiogenesis and endothelial lineage specification, consistent with established ERG functions (*36, 45, 46*) (**Fig. 5C**). By contrast, ERG^WT^ specifically regulated an independent program associated with phosphate metabolism and protein kinase activity (**Fig. 5C**). These findings demonstrate that nucleosome-embedded motif recognition allows a single TF to partition and control distinct biological programs beyond those captured by canonical motif recognition alone.

To test the physiological relevance of this nucleosome-dependent recognition mechanism in an intact organism, we introduced the corresponding mutation into the *Drosophila* ERG ortholog, *Ets21C*, generating transgenic and knock-in flies carrying the mutant *Ets21C*^KKW^ allele (**fig. 9** and **table S8**). Ets21C is a conserved regulator of Jun N-terminal kinase (JNK) signaling, a pathway required for tissue regeneration and stress responses in *Drosophila* (*47–50*). While *Ets21C*^KKW^ flies were viable and fertile, they displayed markedly impaired wing regeneration following genetic tissue ablation compared with wild-type controls (**Fig. 5, D** and **E**, **fig. S9** and **table S8**). Furthermore, *Ets21C*^KKW^ animals exhibited increased sensitivity to oxidative stress, with reduced survival upon paraquat exposure relative to wild-type flies (p < 8.05 × 10^−13^, **Fig. 5F** and **fig. S9**).

Together, these *in vivo* findings elevate nucleosome-derived novel motifs from alternative *in vitro* binding sequences to essential, evolutionarily conserved functional entities that dictate chromatin remodeling, selective gene expression and complex physiological networks beyond the scope of canonical DNA recognition.

## Discussion

Our study shows that nucleosomes do more than restrict TF access to DNA—they reshape TF sequence recognition by creating an additional layer of binding specificity that emerges exclusively within the chromatin environment. By systematically defining the nucleosomal binding preferences of 269 human TFs, we found that many TFs recognize sequence motifs that diverge markedly from their canonical free-DNA counterparts. These nucleosome-dependent motifs are not rare exceptions but are widespread across TF families, contribute causally to chromatin accessibility *in vivo*, and are preferentially enriched in cell-type-specific regulatory elements. Together, these findings indicate that TF binding specificity cannot be fully understood from naked DNA alone, but instead reflects a dynamic interplay between primary DNA sequence and nucleosome architecture.

Several fundamental features distinguish nucleosome-dependent recognition from canonical TF binding. Although members of the same structural family generally recognize highly similar motifs on free DNA, nucleosome-derived motifs frequently diverge among individual family members, revealing substantially greater functional diversification within TF families than is apparent from canonical motifs alone. Moreover, whereas canonical binding preferences remain largely independent of genomic position, nucleosome-dependent recognition is strictly constrained by both the translational and rotational setting of the binding site on the nucleosome. Structural analyses further suggest that these altered specificities arise from mechanisms distinct from those underlying canonical motifs on naked DNA. Rather than being explained solely by intrinsic protein–DNA readout, nucleosome-derived motifs reflect the unique structural environment created by nucleosomal DNA deformation coupled with direct histone contacts. These observations firmly establish the nucleosome as an active determinant of TF sequence recognition, extending its biological role far beyond a simple barrier to binding.

Our results further suggest that nucleosome-derived motifs represent a distinct regulatory and evolutionary layer of gene control. Compared with canonical motifs, they display lower evolutionary sequence conservation (p < 6 × 10^−12^, **fig. S8** and **table S9**) and are preferentially associated with nucleosome-occupied regulatory DNA, consistent with the idea that they encode structural features of chromatin rather than primary DNA sequence alone. This organization provides an elegant mechanism for generating regulatory specificity, allowing closely related TFs to acquire distinct chromatin-binding repertoires without altering their canonical sequence preferences. Such context-dependent recognition helps explain how TFs with highly similar DNA-binding domains achieve non-redundant functions during development and cell differentiation—a notion punctuated by our *in vivo* demonstration that disrupting this nucleosome-specific recognition mode selectively impairs essential, evolutionarily conserved physiological processes such as tissue regeneration.

More broadly, these findings extend the prevailing view of TF specificity from a one-dimensional DNA sequence code to a multidimensional, chromatin-dependent recognition framework in which DNA sequence and nucleosome architecture jointly determine TF binding. This framework provides a mechanistic basis for interpreting TF occupancy in chromatin, refining models of *cis*-regulatory sequence function, and understanding how cell-type-specific gene expression programs emerge from a common genome. The comprehensive resource of nucleosome-binding specificities generated here provides a foundational cornerstone for future efforts to model TF binding *in vivo* and to decode the multi-layered regulatory information embedded within chromatin.

## Acknowledgments

We thank Xiaojing Li, Wei Li and Ying Yang for technical assistance, Fangjie Zhu for help with the NCAP-SELEX technique, and Yihan Chen and Yanhui Xu for comments on the manuscript and insightful discussions. We also thank Mirka Uhlirova from the University of Cologne for the *Drosophila melanogaster* w¹¹¹; Ets21C^Δ10^ line.

We thank the staff at the cryo-EM centers of the National Facility for Protein Science (Shanghai Zhangjiang Lab) and the CAS Center for Excellence in Molecular Plant Sciences for cryo-EM facility access and technical support during data collection. We also thank the Core Facility of Drosophila Resource and Technology (CAS Center for Excellence in Molecular Cell Science) for generating transgenic and knock-in flies, and the Instrument Analysis Center of Tongji University and Core Facilities at Tongji University School of Medicine for data collection and computational resources.

## Author contributions

Y.Y., K.X., and L.W. conceived and designed the experiments. X.Z. performed the NCAP-SELEX experiments. Y.T., Y.H., and D.Z. performed cryo-electron microscopy experiments. Y.T. and L.Z. solved the cryo-EM structures. J.L., X.Y., Y.H., and X.Z. contributed to the generation of doxycycline-inducible TF-expressing hES cell lines and performed ChIP-seq, ATAC-seq, and RNA-seq experiments. H.S. designed the constructs for embryo injection and performed genetic ablation and lifespan assays. Z.X. and Y.Y. developed computational pipelines and performed data analysis. Y.Y. and X.Z. performed the expert motif analysis. Y.Y., K.X., Z.X., and Y.T. prepared the figures and illustrations. Y.Y. wrote the manuscript with input from K.X., L.W., Z.X., X.Z., X.Y., J.L., Y.T., and H.S. All authors contributed to data interpretation and reviewed the final manuscript.

## Data, code, and materials availability

Sequencing data (including ChIP seq, MNase ChIP seq, ATAC seq, RNA seq, and NCAP SELEX) have been deposited in the European Nucleotide Archive (ENA) under accession number PRJEB112530. Structure coordinates for the ERG–nucleosome complexes have been deposited in the Protein Data Bank (PDB) under accession codes 26VU and 26VT. The source code used for data analysis and figure generation is publicly available on GitHub: https://github.com/YinLabTJ/E-MI_computation. All other relevant data are available from the corresponding authors upon reasonable request.

## Materials and Methods

### Cloning, Protein Expression, and Purification

A bacterial protein expression vector incorporating an N-terminal Thioredoxin-6×His tag and a C-terminal 3×FLAG tag was constructed using the pETG20A plasmid as a backbone. The inserts for protein expression were generated as described in Yin *et al*. (*25*) (see **table S1** for protein sequences and domain architectures). Protein expression and purification from *Escherichia coli* (*E. coli*) cells were performed as described in Xie *et al*. (*5*). Briefly, the Rosetta 2(DE3) pLysS strain of *E. coli* (Sigma-Aldrich) was used to express full-length TFs and eDBDs cloned into the pETG20A-6×His-3×FLAG vector. Transformed cells were first cultured overnight in 1 mL of LB medium at 37 °C in 96-well plates (Thermo Fisher Scientific), and then transferred into ZYP5052 auto-induction medium (1:40 dilution, Vincentelli *et al*. (*51*)) supplemented with 0.05% glucose and 0.2% lactose. The cells were then cultured at 37 °C for 8 hours (h), followed by 24 h at 16 °C.

Cells were harvested by centrifugation (4,000 rpm for 15 min) and resuspended in buffer A (300 mM NaCl, 10 mM imidazole in Tris-Cl, pH 7.5) containing 0.5 mg/mL lysozyme (Sigma-Aldrich) and 1 mM PMSF (Sigma-Aldrich). The cells were lysed by one freeze–thaw cycle. DNase I (Sangon Biotech) and MgSO□ were added to the thawed lysate to final concentrations of 10 µg/mL and 1 mM, respectively, to digest released genomic DNA. The lysates were first incubated with Ni–Sepharose 6 Fast Flow resin (Cytiva) for 1 hour on a Timix microplate shaker (Edmund Bühler GmbH) at 1,000 rpm, and then transferred into individual wells of a 96-well Nunc™ filter plate (Thermo Fisher Scientific). Ni–Sepharose beads were washed three times, each with 600 µL of buffer A containing 10 mM and 50 mM imidazole, using a vacuum manifold. Bound proteins were eluted with 500 mM imidazole in buffer A. Purified proteins were assessed by UV absorbance at 280 nm and by SDS–PAGE followed by Coomassie Brilliant Blue staining. Glycerol was added to a final concentration of 50% (v/v) before storage at −20 °C.

The activities of the purified proteins were assessed by HT-SELEX; only those that robustly enriched the expected target sequences were selected for the subsequent NCAP-SELEX process. For selected clones, scale-up cultures (50 or 100 mL) were grown, and proteins were purified using a similar protocol with proportionally increased lysis and wash volumes. The concentration of each purified protein was measured, and when possible, diluted to 50 ng/µL, supplemented with glycerol to a final concentration of 5% (v/v), and stored at −80 °C. ERG proteins (residues 96–418) for cryo-EM sample preparation were expressed in 2–4 L cultures of *E. coli*. Cells were harvested by centrifugation, resuspended in lysis buffer (20 mM Tris–HCl, pH 8.0, 500 mM NaCl), and lysed by two passes through a French press at 15,000 psi. ERG proteins were purified using Ni² –NTA affinity columns (QIAGEN) and eluted with lysis buffer containing 250 mM imidazole. The eluted fractions were pooled and loaded onto a size-exclusion chromatography column (Superdex 200 10/300 GL, GE Healthcare). The peak corresponding to ERG proteins was collected, analyzed by SDS–PAGE, and used for cryo-EM sample preparation.

### NCAP-SELEX Assay

NCAP-SELEX was designed and performed as described previously (*26, 27*) with minor modifications. Briefly, DNA ligands containing either a single 101-bp random sequence or two 30-bp random sequences separated by a truncated 87-bp Widom 601 sequence were designed and used for NCAP-SELEX selection (**table S1**). A 24-bp Illumina sequencing adapter was added to the left side of the ligands, and a 22-bp adapter to the right side, yielding final ligand lengths of 147 bp (lig147) and 193 bp (lig193). Single-stranded oligonucleotides (ssDNA) were purchased from Integrated DNA Technologies (IDT, Ultramer™ DNA Oligos), and double-stranded DNA (dsDNA) ligands were generated by primer extension from the single-stranded templates. Unamplified dsDNA was used as the initial input NCAP-SELEX library, in contrast to our previous study (*26*), in which PCR-amplified dsDNA served as the initial library. Thus, the initial library in this study is substantially more complex (by several orders of magnitude) than that used previously.

The dsDNA ligands were assembled onto histone octamers containing streptavidin-binding protein (SBP)-tagged H2A proteins by stepwise reduction of salt concentration. Specifically, 300 ng of ligands were mixed with H2A-tagged octamers at a 1:2 molar ratio in 5 µL of 2 M NaCl solution. After 30 min incubation at room temperature, the mixture was diluted by sequential addition of TE buffer (2.5 µL, 2.5 µL, 2.5 µL, 2.5 µL, 2.5 µL, 32.5 µL, and 50 µL) supplemented with 1 mM dithiothreitol and 1× cOmplete™ EDTA-free protease inhibitor cocktail (Sigma-Aldrich), with a 1-h incubation after each addition.

After reconstitution, 1.2 µL of streptavidin-coated magnetic Sepharose beads (Cytiva, pre-blocked with 0.5% BSA and 0.1% Tween 20 in 1× incubation buffer) were added to the nucleosomes, and the mixture was incubated at room temperature for 30 min on a Timix microplate shaker at 1,200 rpm. The beads were washed 16 times with 2.28 mL of wash buffer (50 mM KCl, 5 mM NaCl, 1 mM MgCl, 3 µM ZnSO, 100 µM EGTA, 0.1% Tween 20, 1 mM dithiothreitol, 10 mM Tris–HCl, pH 8.0) using a BioTek 405 LRS microplate washer. The total wash buffer volume per sample was twice that used in our previous study¹². Bound nucleosomes were then eluted with 10 mM biotin (Sigma-Aldrich) and 2.5 mM dithiothreitol in incubation buffer (50 mM KCl, 5 mM NaCl, 2 mM MgSO, 3 µM ZnSO, 1 mM K HPO, 20 mM HEPES, pH 7.0, and 100 µM EGTA). Then, 100–200 ng of 6×His-tagged TF protein was added to the eluate, and the mixture was incubated for 30 min at room temperature. After that, 1.8 µL of His Mag Sepharose Ni beads (Cytiva, pre-blocked with 0.5% BSA and 0.1% Tween 20 in 1× incubation buffer) were added. After another 30-min incubation on the Timix shaker at room temperature, the beads were washed 10 times with wash buffer and resuspended in buffer containing 1 mM MgCl and 0.1% Tween 20 in 10 mM Tris–HCl, pH 8.0. DNA was then eluted from the beads by incubation at 90 °C for 8 min. A 12.5-µL aliquot of the bead suspension was used for PCR amplification (25 cycles: 97 °C for 15 s for denaturation, 64 °C for 10 s for annealing, and 72 °C for 36 s for elongation). The amplified ligands were then subjected to additional rounds of NCAP-SELEX selection, and the process was repeated three times. The PCR amplicons from cycle 3 and the initial library were sequenced on an Illumina NovaSeq 6000 or a MGI DNBSEQ-T7 platform using 150-bp paired-end settings.

Prior to NCAP-SELEX, the activities of the expressed TF proteins were assessed by HT-SELEX using lig147 as described in Yin *et al*. (*25*). Only TF proteins that successfully enriched the expected target sequences were included in the NCAP-SELEX experiments. Both reactions were performed in buffers containing 50 mM monovalent cations to reduce nonspecific binding of TF proteins and nucleosomes to the Sepharose beads. The binding of TFs to DNA and to nucleosomes was robust to changes in salt concentration (*25, 26*). The amount of input DNA was sufficient to cover most consecutive and gapped 20-bp sequences, enabling the identification of most, if not all, human TF binding motifs, whose information content is on the order of ∼15 bits (*23, 25*). Additional details of the NCAP-SELEX method and data analysis are available in Zhu *et al*. (*26*).

### Identification of TF Binding Signals on SELEX Ligands

The preferred binding positions of TFs on SELEX ligands were initially analyzed using an unsupervised mutual information (MI)-based approach, which captures binding signals without predefined assumptions (i.e., not restricted to known motifs) (*26*). The underlying rationale is that if a TF contacts nucleosomes, the continuous or spaced 3-mers targeted by the TF will become enriched in SELEX libraries, resulting in increased MI between two non-overlapping positions that accommodate these 3-mers.

MI was initially calculated for all possible 3-mer pairs at two non-overlapping positions on the SELEX ligands, after converting the 3-mer counts to probabilities (i.e., the number of reads containing a specific 3-mer at a given position divided by the total number of reads):

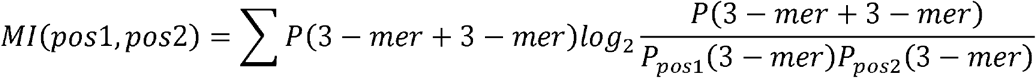

Here, *P*(3-mer+3-mer) denotes the observed probability of a 3-mer pair at position 1 and position 2; *P_pos1_*(3-mer) and *P_pos2_* (3-mer) denote the marginal probabilities of that individual 3-mers at positions 1 and 2, respectively; and MI(pos1, pos2) are the sum of mutual information over all possible 3-mer pairs.

Histones contact the DNA backbone at ∼10-bp intervals and slightly prefer certain DNA sequences over others, causing increased MI between positions separated by 10 bp (*7, 26, 52*). To separate TF-introduced signals from nucleosome-introduced signals, we restricted our MI analysis to the 10 most highly enriched 3-mer pairs (E-MI) in the SELEX libraries, which typically represent the subsequences targeted by TFs. For TF signal analysis on lig147, fragments of 92–109 bp in length were used, and were adjusted to a uniform length of 101 bp by either adding N-padding or trimming extra bases from both ends. For analysis of TF signals on lig193, the 87-bp Widom 601 sequence was converted to Ns to avoid spurious signals. 3-mer pairs that were commonly enriched across SELEX libraries of multiple TFs were considered as noise and masked in downstream analyses. We then focused our analysis on the E-MI diagonals (**table S3**), which represent the TF footprints on the SELEX ligands.

Hierarchical clustering of the E-MI diagonals was performed using the *hclust* function in R with cosine distance and Ward’s D2 linkage criterion. The E-MI diagonals were first log_10_-transformed and then individually normalized by subtracting their means. Circular representations of the clustered E-MI diagonals were visualized using the R package *circlize* (*53*).

### PWM Generation and Similarity Analysis

To model TF binding, we generated position weight matrices (PWMs) for the studied TFs using our previously described Autoseed algorithm (*29*). Briefly, subsequences of 8 bp and 10 bp in length (either gapless or with a central gap) were counted in each SELEX library, and their similarity was assessed using the Huddinge distance measure (*29*). Subsequences with higher counts than any other sequence within a Huddinge distance of 1 were defined as local-maximum k-mers and used as initial seeds for each TF. These seeds were then used to generate initial PWMs for each TF using a multinomial model (*23, 54*). In this multinomial model, the counts of each seed (consensus sequence) and three additional sequences with base substitutions at position *i* were used to generate the PWM at position *i*. This counting process was iterated from position 1 to position *j* (where *j* is the seed length), and the counts at each position were background-corrected and used to generate the initial PWM for each TF (*5, 23, 54*).

Seeds were subsequently refined by: (1) incorporating IUPAC degeneracy codes at positions where the frequency of the most common base was < 0.5, to accommodate more sequence variants; (2) extending to flanking positions when the ratio between the most and least frequent bases at the flanking position was greater than 2; and (3) allowing one mismatch (multinomial parameter = 2) at positions other than the one used for mononucleotide distribution measurement, for seeds longer than 10 bp (*5*). The exact seed sequences and refined multinomial models are listed in **table S2**. The sequence reads have been deposited in the ENA under accession number **PRJEB112530**.

In this study, 996 PWM models were generated for 312 TF proteins from NCAP-SELEX ligands. This motif collection was further expanded by including 1,758 previously published HT-SELEX motifs for human and mouse TFs. Specifically, we collected the following published HT-SELEX motif sets: 707 motifs for 381 human TFs and 134 motifs for 82 mouse TFs from Jolma *et al*. (*23*) (excluding replicate motifs within that study); 31 motifs for 26 human TFs from Jolma *et al*. (*55*) (excluding CAP-SELEX motifs); 10 motifs for 8 human TFs from Nitta *et al*. (*29*) (excluding *Drosophila* TF motifs); 864 motifs for 542 human TFs from Yin *et al*. (*25*) (excluding methyl-SELEX motifs); and 12 motifs for 10 human TFs from Xie *et al*. (*5*) (excluding CAP-SELEX motifs).

Similarities between all pairs of motifs were calculated using TOMTOM v5.3.3 (with a significance threshold of 1) (*56*) and SSTAT (Motif Statistics Software Suite v1.1) (*57*), with the following settings for SSTAT: a 50% GC-content background model, pseudocount regularization, and a stringent type I error threshold of 0.01 to limit the influence of low-affinity sites. The similarity scores between NCAP-SELEX and HT-SELEX motifs were compared with those between HT-SELEX motifs from this study and those from previous studies. NCAP-SELEX motifs with similarity scores < 5×10^−6^ (SSTAT Ssum) or Q-values > 0.001 (TOMTOM) to any HT-SELEX motif were classified as novel. This analysis yielded 268 novel motifs.

A motif network was constructed, in which nodes representing TF binding models were connected if their SSTAT Ssum similarity score exceeded 1.5×10^−5^. The network was visualized using Cytoscape v3.10.1 (*58*). The minimum dominating set of the network was defined as the smallest set of PWMs that are connected to every PWM outside the set. Dominating set analysis of the 2,754-motif collection was performed using GLPSOL (GLPK LP/MIP solver) v4.65, with solver configurations following the settings described in Jolma *et al*. (*55*). This analysis uncovered 669 representative motifs with distinct sequence specificities. Of these 268 novel motifs, 142 were identified as representative motifs (**table S2**).

### Motif Matching, Enrichment and Occupancy Analysis

The NCAP-SELEX motifs were matched to their original NCAP-SELEX ligands (from which they were recovered) using MOODS v1.9.4.1 (*28*) with a P-value threshold of 0.0001. The resulting motif matches were used to complement the systematic E-MI analysis of TF binding positions on SELEX ligands.

The 268 novel NCAP-SELEX motifs and their 179 most closely related HT-SELEX counterparts (referred to as canonical motifs) were also matched to the repeat-masked human genome (GRCh38) using MOODS v1.9.4.1 with a P-value cut-off of 0.0001 and a score cut-off of 5, to obtain a large set of putative binding sites for each corresponding TF. These motif matches were then used to evaluate whether each motif was enriched or depleted at cell-type-specific candidate *cis*-regulatory elements (cCREs). The cCREs were derived from 111 human cell types in the *cis*-element atlas (*34*) (CATlas; https://catlas.org/humanenhancer; downloaded on July 14, 2025). In CATlas, cCREs are defined as non-coding chromatin regions that are accessible to nucleases and were identified by single-cell ATAC-seq across 30 adult human tissues. In total, approximately 400,000 cCREs were found to exhibit cell-type-restricted ATAC-seq signals and were classified into 15 cell-type-specific groups.

For each biosample, we computed the fraction of cell-type-specific cCREs that overlapped with matches of each motif model. To assess cell-selective enrichment, we calculated the median and standard deviation of the fractional occurrence of each motif across all biosamples. Outliers exceeding 3.5 times the median absolute deviation (MAD) were removed, and the remaining values were log1p-transformed and used to parametrize a normal distribution. From the fitted normal distribution, an upper-tail P-value was derived for each PWM model in every biosample (**table S5**). The P-values for each motif across all biosamples were subjected to false discovery rate (FDR) correction using the Benjamini–Hochberg method. The corrected P-values were defined as the enrichment scores of the motif in each tested biosample.

To further evaluate TF occupancy at motif matches within open chromatin regions, we downloaded per-nucleotide footprint statistics files for 243 biosamples from the ENCODE database (*44*) (https://resources.altius.org/~jvierstra/projects/footprinting.2020; downloaded on June 6, 2025). A motif match was considered to be occupied by a TF if the matched site overlapped a DNase I hypersensitive site (DHS; listed in the peaks.bed.starch files at the same URL) and the FPR-adjusted P-values of all nucleotides within the site were < 0.05. Nucleotides within the motif matches that were located at positions with low information content (< 0.3 bits) in the PWM model were excluded from the analysis. The fractions of motif match sites occupied by TFs were computed for each motif in each biosample (**table S7**). These fractions were compared between each novel motif and its related HT-SELEX counterpart in three biosamples — the ones in which the HT-SELEX motifs showed the highest enrichment at DHSs across the dataset.

### Analysis of Motif Contributions to Chromatin Accessibility

To identify causal sequence features influencing chromatin accessibility, we trained ChromBPNet models — supervised convolutional neural networks — to predict the accessibility profile and the total natural-log-transformed counts in the central windows of the input regions, following the workflows described in previous studies (*38, 39*).

To prepare genomic regions for training, we downloaded ATAC-seq peak and fragment files for each of the 110 human cell types in CATlas (*34*) (https://catlas.org/humanenhancer; downloaded on July 14, 2025). The acinar cell type was excluded from the analysis due to an incomplete peaks.bed file. For each cell type, the 2,114-bp DNA sequences flanking the summits of the top 30,000 ATAC-seq peaks were used for downstream analysis.

We followed the workflow described in the ChromBPNet package (https://github.com/kundajelab/chrombpnet; version 1.0.1) to train the models. First, the *chrombpnet prep nonpeaks* command was run to define background regions devoid of ATAC-seq peaks but with GC content similar to that of the peak regions. Second, the *chrombpnet bias pipeline* command was run to learn the enzymatic bias in the ATAC-seq assay using fold 0 (the first fold of the five-fold cross-validation scheme) of the cardiomyocyte cell type in CATlas, with the bias threshold factor set to -b 0.4. After interpreting the bias model using DeepLIFT (*40*), we confirmed that it had learned six of the eight Tn5 motifs but none of the TF motifs. Third, the *chrombpnet pipeline* command, together with the trained bias model, was used to train ChromBPNet models for five folds for each of the 110 cell types in CATlas, yielding 550 models in total.

All models were trained on the ENCODE GRCh38 reference genome (GRCh38_no_alt_analysis_set_GCA_000001405.15) using chromosome size information obtained from ENCODE under accession number ENCFF667IGK. Each fold (0–4) contained a different partition of chromosomes into training, validation, and test sets, and each chromosome was assigned to the test set in at least one fold. The default human chromosome folds provided with ChromBPNet were used (*38, 39*). Each model was assessed based on the Pearson and Spearman correlations between the predicted and observed log counts in peaks located on held-out test chromosomes. Models for five cell types were excluded from downstream analysis due to low Spearman correlation (< 0.5; **table S6**).

Finally, the *chrombpnet interpret* command was run to apply the DeepLIFT algorithm (*40*) and compute contribution scores for each nucleotide in the 2,114-bp input windows with respect to the predicted counts. Contribution scores were derived for each model fold across all peak regions and then averaged across folds. The averaged predicted accessibility profiles and contribution scores were used for downstream analyses.

We next investigated the extent to which the novel and corresponding canonical motifs were predictive of accessibility in a given cell type, by comparing the contribution of their motif matches to that of artificial control motif matches. To generate a set of artificial control motifs, each novel or canonical motif was first split into two partial motifs at all possible cut positions. Control motifs were then constructed by concatenating each pair of partial motifs in three alternative orientations: (i) right half + left half; (ii) left half + right-half reverse complement; and (iii) left-half reverse complement + right half. The resulting artificial PWMs were compared to the original PWM using *motifsimilarity* (https://github.com/jutaipal/motifsimilarity), and those with similarity scores < 0.1 were retained as artificial control motifs. The control motifs were also matched to the repeat-masked human genome (GRCh38) using MOODS. For each motif match, the contribution scores of the nucleotides within it were averaged to obtain a single contribution score per match. The averaged contribution scores of all matches of a given motif in a cell type were collected. The score distributions were compared between each novel or canonical motif and its corresponding control motifs in each cell type using a one-sided Wilcoxon test. P-values were adjusted using the Benjamini–Hochberg (BH) method. A motif was considered predictive of accessibility in a cell type if the adjusted P-value was < 0.05 and the mean score across the collection was > 0.002.

We also performed de novo motif discovery on sequences driving chromatin accessibility and subsequently measured the similarity of the identified motifs to the novel and canonical motifs. Briefly, TF-MoDISco (https://github.com/kundajelab/tfmodisco/tree/v0.5.6.5) was first used to identify short contiguous stretches of nucleotides with high positive or negative importance scores (referred to as “seqlets”), which were then clustered into recurrent 30-bp patterns. A total of 50,000 seqlets were then sampled from the central 400 bp of the input regions. *De novo* motifs were derived from these seqlets, with each motif represented by a contribution weight matrix (CWM; 4×30) and a position probability matrix (PPM). CWMs learned from three of the 105 cell types in CATlas were found to exhibit low complexity; ChromBPNet models and motifs for these three cell types were excluded from downstream analysis. In total, 13,177 motifs were learned from 102 cell types (510 models). Similarities between these motifs and the novel or HT-SELEX motifs were calculated using TOMTOM v5.3.3 and SSTAT (Motif Statistics Software Suite v1.1). Of the 13,177 de novo motifs, 1,300 were found to be more similar to the novel motifs than to the HT-SELEX motifs. Representative examples are shown in **fig. S8.**

### Evolutionary Conservation Analysis

To measure the conservation of TF–nucleosome interactions reflected by the motifs, we followed the procedure described in Xie *et al*. (*5*). Briefly, we used the conservation scores and constrained nucleotides defined by the Zoonomia Consortium (*59, 60*) to assess the potential conservation of genomic sites recognized by the novel motifs, the canonical motifs, and their artificial control motifs (constructed as described above).

By comparing genome sequences of 241 placental mammals, the Zoonomia Consortium assessed each base in the human genome and classified it as either slower than expected (conservation) or faster than expected (acceleration) based on phyloP scores (*61, 62*) (positive scores indicate conservation; negative scores indicate acceleration). The most significantly conserved nucleotides (phyloP score ≥ 2.27 and FDR < 0.05; covering ∼101 Mb, or 3.26% of the genome) were defined as the human constrained nucleotides. Constrained nucleotides overlapping protein-coding exons (GENCODE v44) or simple tandem repeats (UCSC) were excluded from downstream analysis.

For each true motif (i.e., novel or canonical motif), average phyloP scores were calculated for its individual matches that overlapped with at least one constrained nucleotide. A two-component Gaussian mixture model (GMM) was fitted to these scores to separate constrained matches from non-constrained ones. Ten sets of initial parameters were tested for GMM fitting, and the best model was selected based on the Bayesian information criterion (BIC) for further analysis. From the selected model, we derived a threshold value for each motif, defined as the score at which the posterior probability of belonging to the constrained component exceeded that of the non-constrained component. Thus, a motif match with an average phyloP score above this threshold was classified as conserved. Different thresholds were obtained for each motif (**table S9**).

To assess the evolutionary conservation of the novel motifs, we first selected 10,000 non-overlapping matches for each true motif and its corresponding control motifs, based on MOODS scores. Average phyloP scores were then computed for each of these matches. Matches with average phyloP scores exceeding the motif-specific threshold were assigned to the conserved set; the remaining matches were assigned to the non-conserved set. For each true motif, we recorded: (i) the number of conserved (k) and non-conserved (n − k) matches among its own top 10,000 matches; and (ii) the number of conserved (m − k) and non-conserved (N + k − n − m) sites among the top 10,000 matches of its corresponding control motifs, where n is the total number of true motif matches, N is the total number of motif matches, and m is the total number of conserved sites among matches. A P-value was then computed using a hypergeometric test to assess whether conserved sites were enriched among the matches of each true motif compared to its control motifs. The P-values for all true motifs were adjusted to control the family-wise error rate (FWER) using Holm’s method (*5*). Motifs with FWER-adjusted P-values < 0.05 were defined as conserved. Full statistics are provided in **table S9**.

To determine whether the novel NCAP-SELEX motifs were more conserved than their related HT-SELEX counterparts, we computed the numbers of conserved and non-conserved motifs for both sets and applied Fisher’s exact test to evaluate the statistical significance. The enrichment fold for each true motif was calculated as (k/n) / ((m − k)/(N − n)) and plotted against the number of conserved matches among the 10,000 highest-scoring matches (**fig. S8**).

### Histone Octamer and DNA Preparation

Full-length histone sequences from *Homo sapiens* were cloned into pETDuet and pCDFDuet expression vectors as described in Xue *et al*. (*27*). H2A and H3 were each fused at the N-terminus to a 6×His–SUMO tag, with a PreScission protease cleavage site inserted between the histone and the SUMO tag. Histone proteins were expressed in *E. coli* BL21(DE3) cells and induced with 0.5 mM IPTG. Histone proteins were purified using Ni² –NTA affinity columns (QIAGEN) and eluted with buffer containing 20 mM Tris–HCl (pH 8.0), 2 M NaCl, 250 mM imidazole, and 10% (v/v) glycerol. After cleavage of the His–SUMO tag, H2A–H2B and H3–H4 dimers were mixed at a 1:1 molar ratio and incubated overnight at 4 °C. The sample was concentrated and loaded onto a size-exclusion chromatography column (Superdex 200 10/300 GL, Cytiva). The peak corresponding to the intact octamer was collected, analyzed by SDS–PAGE, and used for nucleosome reconstitution. For NCAP-SELEX assays, histone octamers were prepared following the same protocol, except that an additional streptavidin-binding peptide (SBP) tag was inserted between the PreScission cleavage site and H2A.

For nucleosome reconstitution, two DNA templates (template-1 and template-2) were derived from ERG NCAP-SELEX experiments. To eliminate potential ERG binding, two GGAA sites in the adapter regions of both templates were mutated to GCTT, yielding template-1a and template-2a. The full sequences of all four templates are provided in **table S1**. DNA templates 1a and 2a were synthesized by GenScript Biotech and amplified by PCR. PCR products were purified on a HiTrap Q HP column (Cytiva) using an ÄKTA FPLC system, with DNA fragments eluted by a linear salt gradient from 0.15 M to 1 M NaCl. Peak fractions were analyzed by agarose gel electrophoresis, and those containing the expected DNA fragments were pooled. The pooled DNA was concentrated by ethanol precipitation and stored at −20 °C until use.

### Nucleosome Reconstitution

Nucleosomes were reconstituted from histone octamers and DNA templates using a salt-gradient dialysis method, as previously described by Dyer *et al*. (*63*). Briefly, histone octamers and DNA were mixed at a 1.4:1 molar ratio in high-salt buffer (2 M NaCl, 20 mM HEPES, pH 7.5, 1 mM EDTA) and transferred into SnakeSkin™ dialysis tubing (Thermo Fisher Scientific). The dialysis tubing containing 1 mL of the mixture was placed in 450 mL of high-salt buffer, into which 4.5 L of no-salt buffer (20 mM HEPES, pH 7.5, 1 mM EDTA) was gradually pumped at a flow rate of 3 mL/min over 20 hours. This resulted in a gradual reduction of salt concentration from 2 M to approximately 180 mM. The dialysis tubing was then transferred to 500 mL of fresh no-salt buffer and incubated for an additional 4 hours to ensure complete removal of residual salt. The reconstituted nucleosomes were analyzed by electrophoresis on a 6% native polyacrylamide gel.

### Cryo-EM Sample Preparation and Data Collection

ERG protein was added at a 5-fold molar excess to reconstituted nucleosomes (1 μM, assembled with either template-1a or template-2a DNA) in a buffer containing 20 mM HEPES (pH 7.5), 1 mM EDTA, 100 mM NaCl, and 1 mM DTT. The mixtures were incubated on ice for 30 min and then concentrated using 0.5 mL Amicon Ultra centrifugal filters (100 kDa MWCO; Sigma-Aldrich) to a final nucleosome concentration of approximately 5–10 μM, as determined by absorbance at 260 nm. The resulting samples were then applied to cryo-EM grids following the procedure described in Xue *et al*. (*27*). Briefly, Quantifoil® R 1.2/1.3 300-mesh gold grids were glow-discharged using a SOLARUS plasma cleaning system for 135 s; 3.5 µL of each sample was applied to a grid in a Vitrobot Mark IV (Thermo Fisher Scientific) at 4 °C and 100% humidity, and the grids were immediately vitrified by plunging into liquid ethane.

For the ERG–nucleosome complex assembled with template-1a (referred to as ERG–Nuc^ACCGGT^), data were collected automatically using EPU 3.15 (Thermo Fisher Scientific) on a 300 kV Krios G4 electron microscope equipped with a Falcon 4 direct electron detector. Images were recorded at a nominal magnification of 96,000× with a pixel size of 0.83 Å. The exposure time was set to 5.54 s, with a total accumulated dose of 55 e /Å². Images were collected with a defocus range of −1.0 to −1.8 μm.

For the ERG–nucleosome complex assembled with template-2a (referred to as ERG–Nuc^AGGAAG^), cryo-EM images were acquired using EPU 2.12 (Thermo Fisher Scientific) on an FEI Titan Krios operating at 300 kV, with a nominal magnification of 81,000×. Images were recorded using a K3 Summit direct detector (Gatan) in super-resolution mode, with a pixel size of 0.89 Å. The defocus range was set to −1.0 to −1.6 μm. Thirty frames were recorded per stack, with an exposure time of 2.17 s and a total dose of 55 e /Å².

### Cryo-EM Data Processing and Analysis

Processing details are summarized in **table S4** and **fig. S5** and **S6**. For each dataset, all image stacks were motion-corrected using MotionCor2 (*64*) within RELION 3.1 (*65*), and contrast transfer function (CTF) parameters were estimated using CTFFIND4 (*66*). Particles were automatically picked using the Blob picker in cryoSPARC v5.0.6 (*67*). Initial references for the ERG–nucleosome complexes were generated *ab initio* in cryoSPARC, low-pass filtered to 20 Å, and then used as starting models for 3D classification of all datasets. For each dataset, to expedite computation, particles were extracted after 4× binning (i.e., bin4) of the pixel size and subjected to several rounds of 2D and 3D classification in cryoSPARC. Classes exhibiting high-resolution features were selected for further processing. The selected particles were then re-extracted without binning (i.e., 1× binning, original pixel size) using a box size of 256 pixels, and subjected to another round of 3D classification and cleaning. All datasets were processed without imposing symmetry (*C*1). Final Fourier shell correlation (FSC) curves, along with directional FSC curves and anisotropy estimates, were calculated using the 3DFSC server (**fig. S5** and **S6**). Local resolution for each map was also calculated in cryoSPARC (**fig. S5** and **S6**).

For the ERG–Nuc^ACCGGT^ complex dataset, classes exhibiting additional densities were selected after heterogeneous refinement. Both classes, which displayed a clear additional density at superhelical location (SHL) −1.7, were selected and merged. The selected subset was then subjected to a round of focused classification using a small soft spherical mask centered on the additional density near SHL −1.7. A class showing strong additional density was selected, and further heterogeneous refinement was performed. As a final step, the dataset was subjected to non-uniform refinement in cryoSPARC, which improved the local resolution distribution in the 3D reconstruction. The final map was sharpened with a B-factor of −93.1. The final dataset comprised 170,908 particles.

The ERG–Nuc^AGGAAG^ complex dataset was processed similarly. After initial coarse cleaning, the dataset was classified into four classes, two of which were of high quality. Both classes displayed a clear additional density at SHL −2.3 and were merged. The selected subset was then subjected to a round of focused classification using a small soft spherical mask centered on the additional density near SHL −2.3. As a final step, the dataset was subjected to non-uniform refinement in cryoSPARC, yielding a map of the ERG–Nuc^AGGAAG^ complex that was sharpened with a B-factor of −110.3. The final dataset comprised 77,603 particles.

In both cases, the final reconstruction was obtained from a subset of high-quality particles, yielding a single high-resolution structure for each complex.

### Model Fitting and Refinement

The cryo-EM structures of the nucleosome (PDB: 6T79) and ERG (PDB: 4IRI) were used as starting models. The models were initially fitted into the cryo-EM density maps using UCSF Chimera v1.11.1 (*68*) and then manually adjusted in Coot v0.8.9.2 (*69*). Model optimization was performed through iterative cycles of real-space refinement in PHENIX v1.21.2-541969 (*70*) and manual adjustment in Coot. Secondary structure and Ramachandran restraints were applied during refinement. The final models were validated using MolProbity; model-to-map correlation coefficients were calculated in PHENIX. Statistics for model validation are provided in **table S4**.

### Electrophoretic Mobility Shift Assay (EMSA)

For affinity analysis, 5 µL mixtures containing 1 pM DNA or nucleosome and 0–15 pM purified wild-type or mutant ERG were prepared in either 1× binding buffer (50 mM NaCl, 10 mM Tris, pH 8.0, 1 mM EDTA, pH 8.0, 1 mM DTT) or 1× incubation buffer (20 mM HEPES, pH 7.0, 50 mM KCl, 5 mM NaCl, 2 mM MgSO, 1 mM K HPO, 3 µM ZnSO, 100 µM EGTA, 2.5 mM DTT, 10 mM biotin). The DNA sequence (GTCTCGTGGGCTCGGACCGGAAGTGG) contained a canonical ERG binding site (ACCGGAAGT). Specifically, nucleosomes assembled with template-1a (**table S1**) were incubated with wild-type ERG or the ERG^KKW^ mutant; nucleosomes assembled with template-2a (**table S1**) were incubated with wild-type ERG or the ERG^IDR–/–^ mutant; and free DNA was incubated with each ERG protein separately. The mixtures were incubated at room temperature for 1 h in protein LoBind tubes (Eppendorf), then mixed with Novex™ Hi-Density TBE sample buffer (Thermo Fisher Scientific), and loaded onto 6% (for nucleosome–ERG complexes) or 12% (for DNA–ERG complexes) non-denaturing polyacrylamide gels. Electrophoresis was performed in 0.5× TBE buffer at 200 V for 30 min (for nucleosome–ERG samples) or 40 min (for DNA–ERG samples). Gels were stained with SYBR™ Gold dye (Thermo Fisher Scientific), washed, and imaged using a Bio-Rad ChemiDoc MP imaging system. Three independent experiments were performed for each sample.

### Cell Culture

HepG2 cells were obtained from the American Type Culture Collection (ATCC; HB-8065) and cultured at 37 °C in a humidified incubator in Eagle’s Minimum Essential Medium (EMEM; ATCC, 30-2003) supplemented with 10% (v/v) fetal bovine serum (FBS; Thermo Fisher Scientific, 10100147c) and 1% penicillin–streptomycin (Thermo Fisher Scientific, 15140122). VCaP cells (CAS Cell Bank, Beijing, China; cat. no. SCSP-5034) were cultured in DMEM (Thermo Fisher Scientific, 10569010) supplemented with high glucose, GlutaMAX™, sodium pyruvate, 10% FBS, and 1% penicillin–streptomycin. Primary mouse embryonic fibroblasts (MEFs) were isolated from CF-1 mouse embryos at embryonic day 13.5 (E13.5) and cultured in MEF medium consisting of DMEM (Thermo Fisher Scientific, 11965118), 10% FBS, and 1% penicillin–streptomycin. H9 human embryonic stem cells (ESCs) were maintained on irradiated MEF feeder layers in hES cell culture medium consisting of DMEM/F12 (Gibco, 11330-032), 20% knockout serum replacer (Gibco, 10828-028), L-glutamine (Gibco, 25030149), 1× MEM non-essential amino acids (Gibco, 11140-050), 0.1 mM β-mercaptoethanol (Amresco, 0482-100ml), and 4 ng/mL fibroblast growth factor 2 (FGF-2; Peprotech, 100-18B). hES cells were cultured at 37 °C with 5% CO and passaged every 5 days using Dispase (Gibco, 17105-041) digestion.

### Generation of Doxycycline-Inducible TF-Expressing human ES Cell Lines

To generate H9 humsn ES cells capable of doxycycline-inducible TF expression, we first engineered a parental line carrying a CAG–rtTA–IRES–BSD cassette integrated upstream of the EEF1A1 gene promoter (*71*). In this background, the coding sequences of POU3F3, wild-type ERG, or mutant ERG were targeted to the AAVS1 safe-harbor locus.

Coding sequences of POU3F3 (from neural progenitor cells) and wild-type ERG (from VCaP cells) were amplified by RT–PCR from total RNA. The ERG coding sequence corresponds to UniProt accession P11308-3. Reverse PCR primers were designed to append sequences encoding a GSG linker and two tandem HA tags (YPYDVPDYA) to the 3′ ends of the amplified POU3F3 and ERG open reading frames. The ERG coding sequence was further mutagenized to introduce point mutations D350K, D352K, and D366W, generating the ERG^KKW^ mutant. Each coding sequence (POU3F3, wild-type ERG, or mutant ERG) with its C-terminal GSG–2×HA tag was cloned into the TRE-TIGHT-EGFP-backward donor plasmid (Addgene, 22077) by replacing the GFP open reading frame using the ClonExpress Ultra One-Step Cloning Kit (Vazyme), following the manufacturer’s instructions.

The following plasmids were prepared using an endotoxin-free Midi kit (QIAGEN) and quantified with a Qubit 4 fluorometer (Invitrogen): TRE-TIGHT-POU3F3-backward donor, TRE-TIGHT-ERG^WT^-backward donor, TRE-TIGHT-ERG^KKW^-backward donor, and the left and right hAAVS1 TALEN plasmids (Addgene, 52341 and 52342).

TALEN-mediated homologous recombination was performed to insert POU3F3, wild-type ERG, and mutant ERG into the AAVS1 safe-harbor locus, as described by Hockemeyer *et al*. (*72*). The EEF1A1-rtTA H9 human ES cells were cultured in ES cell medium supplemented with 10 µM ROCK inhibitor (Y-27632; Selleck Chemicals) for 24 hours and then dissociated with Accutase (Thermo Fisher Scientific) for 5–6 min. Approximately 1–5 × 10^6^ cells were resuspended in 200 µL of Gene Pulser electroporation buffer (Bio-Rad), and 40 µg of one of the donor plasmids and 5 µg of each of the left and right AAVS1 TALEN plasmids were added. Electroporation was performed using a Gene Pulser Xcell system (Bio-Rad) with settings of 250 V, 500 µF, and 0.4-cm cuvettes. Cells were then plated at low density onto MEF feeder layers in human ES cell culture medium, with 10 µM ROCK inhibitor added during the first 24 hours.

Puromycin selection (0.5 µg/mL) was started 48 h after electroporation and maintained with daily medium changes for 10–14 days. Drug-resistant clones were then manually picked and expanded in the presence of puromycin for an additional 6–7 days. Expanded clones were genotyped by PCR and Sanger sequencing. Site-specific integration at the AAVS1 locus was confirmed by genomic PCR with junction-spanning primers (**table S1**). Clones with confirmed biallelic integration and doxycycline-inducible expression of the corresponding TF protein (as verified by western blot) were used for downstream experiments.

### Chromatin Immunoprecipitation Sequencing (ChIP-seq) and MNase-ChIP-seq

ChIP-seq was performed on the previously established H9 human ES cell lines carrying site-specific integrations of the TRE-TIGHT-POU3F3-backward, TRE-TIGHT-ERG^WT^-backward, or TRE-TIGHT-ERG^KKW^-backward donor constructs, following a published protocol (*73*). Briefly, cells were plated on 6-well plates coated with 0.1% gelatin (Sigma-Aldrich) and cultured on MEF feeder layers in human ES cell culture medium for 4 days until reaching ∼70% confluency. Doxycycline (Sigma-Aldrich) was then added to the culture medium at a final concentration of 1 µg/mL. Cells were cultured for an additional 24 h in human ES cell culture medium containing doxycycline and then harvested by Accutase digestion at 37 °C for 5–6 min. Harvested cells were fixed in 1% formaldehyde (Thermo Fisher Scientific) for 10 min at room temperature, and crosslinking was quenched by adding glycine to a final concentration of 0.125 M. Cells were washed twice with ice-cold PBS, counted using an automated cell counter (Invitrogen), and resuspended in high-salt lysis/sonication buffer (800 mM NaCl, 25 mM Tris–HCl, pH 7.5, 5 mM EDTA, 1% Triton X-100, 0.1% SDS, 0.5% sodium deoxycholate) supplemented with cOmplete™ protease inhibitor cocktail (Roche) to a density of 1 × 10□ cells per 100 µL of buffer. Cells were aliquoted into TPX tubes (Diagenode) at 3 × 10 cells per tube (one sample). Chromatin was sheared using a Diagenode Bioruptor Plus with 30 s ON / 90 s OFF cycles at high power. Sonication was performed for 20 cycles at 4 °C to generate an average fragment size of 100–500 bp. The samples were then centrifuged at 13,000 rpm for 15 min at 4 °C, and the supernatant was collected. Supernatants from cells of the same genotype were pooled and diluted 1:5 with chromatin dilution buffer (25 mM Tris–HCl, pH 7.5, 5 mM EDTA, 1% Triton X-100, 0.1% SDS) containing protease inhibitors. A 50-µL aliquot of the diluted chromatin was stored at −20 °C as the input fraction.

For immunoprecipitation, 20 µL of Dynabeads Protein G magnetic beads (Thermo Fisher Scientific) were coupled with 3 µg of anti-HA antibody (Diagenode) or non-specific IgG (Abcam) in 40 µL of chromatin dilution buffer for 3 h at room temperature with rotation. Approximately 1.2 mL of diluted chromatin was added to the antibody-coated beads for each sample, and the mixture was incubated overnight at 4 °C with rotation (15–20 rpm). After incubation, the beads were washed once with wash buffer A (140 mM NaCl, 50 mM HEPES, pH 7.9, 1 mM EDTA, 1% Triton X-100, 0.1% SDS, 0.1% sodium deoxycholate), once with wash buffer A containing 500 mM NaCl, and once with wash buffer C (250 mM LiCl, 20 mM Tris– HCl, pH 7.5, 1 mM EDTA, 0.5% NP-40, 0.5% sodium deoxycholate). After two washes with TE buffer, the beads were resuspended in 200 µL of elution buffer (1% SDS, 10 mM Tris–HCl, pH 7.5, 1 mM EDTA) and incubated at 65 °C for 5 min, followed by 15 min at room temperature with rotation. NaCl and RNase A (Thermo Fisher Scientific) were added to both the eluted and input samples to final concentrations of 160 mM and 20 µg/mL, respectively. Samples were incubated overnight at 65 °C to reverse crosslinking, followed by addition of 200 µg/mL proteinase K and 5 mM EDTA (Thermo Fisher Scientific) and incubation at 45 °C for 2 h. Purified DNA was obtained by phenol–chloroform extraction, and libraries were prepared using the NEBNext Ultra II DNA Library Prep Kit (NEB) for Illumina sequencing. Paired-end sequencing (2 × 150 bp) was performed on an Illumina NovaSeq X Plus platform.

MNase-ChIP-seq was performed according to a published protocol (*74*). Briefly, 10 × 10^6^ HepG2 cells were harvested and crosslinked in 10 mL of culture medium containing 1% formaldehyde for 10 min at room temperature. Crosslinking was quenched with 0.125 M glycine for 5 min. After two washes with ice-cold DPBS, cells were resuspended in 10 mL of 0.5× PBS containing 0.5% Triton X-100 and incubated on ice for 3 min. Nuclei were pelleted by centrifugation at 500 × g for 4 min at 4 °C and resuspended in 1× MNase digestion buffer containing 100 µg/mL RNase A to a density of 2 × 10□ nuclei per 100 µL of buffer. MNase (50 units; NEB) was added to 2 × 10^6^ nuclei, and the mixture was incubated at 37 °C for 30 min, followed by addition of 100 µL of stop buffer (40 mM EDTA, 40 mM EGTA, 5 mg/mL BSA, 150 mM LiCl, 2 mM TCEP). The samples were centrifuged at 13,000 × g for 1 min, and the supernatant from 1 × 10^7^ nuclei was incubated with 20 µL of Protein G Dynabeads pre-coated with 3 µg of anti-CEBPB antibody (GeneTex) or non-specific IgG (Abcam). DNA fragments were then enriched and libraries were prepared for Illumina sequencing following the same protocol as described above for ChIP-seq.

### ATAC-seq and RNA-seq Profiling of Doxycycline-Inducible ERG ES Cells

ATAC-seq was performed on the previously established H9 human ES cell lines carrying site-specific integrations of the TRE-TIGHT-ERG^WT^-backward or TRE-TIGHT-ERG^KKW^-backward donor constructs, following a published protocol (*75*). Briefly, approximately 2 × 10^6^ cells from each line were harvested before doxycycline treatment, and at 24 h and 48 h after treatment with 1 µg/mL doxycycline. Cells were dissociated by Accutase digestion at 37 °C for 5–6 min. After two washes with ice-cold PBS, cells were resuspended in 3 mL of ice-cold lysis buffer (10 mM Tris–HCl, pH 7.4, 10 mM NaCl, 3 mM MgCl, 0.1% NP-40, 0.1% Tween-20, and 0.01% digitonin) and incubated on ice for 3 min with gentle pipetting.

Nuclei were collected by centrifugation at 500 × g for 5 min at 4 °C. Nuclei were resuspended in 5–10 mL of wash buffer (10 mM Tris–HCl, pH 7.4, 10 mM NaCl, 3 mM MgCl, and 0.1% Tween-20) to a density of 2 × 10^5^ nuclei per mL. Approximately 50,000 nuclei (250 µL) were transferred to a fresh tube, pelleted by centrifugation, and resuspended in 50 µL of tagmentation mix containing 25 µL of 2× Tagment DNA Buffer, 16.5 µL PBS, 0.5 µL of 10% Tween-20, 2.5 µL Tn5 transposase, and 5.5 µL nuclease-free water. Tagmentation was carried out at 37 °C for 30 min in a ThermoMixer with shaking at 1,000 rpm. Tagmented DNA was purified using a MinElute Reaction Cleanup Kit (QIAGEN) following the manufacturer’s instructions and eluted with 22 µL of nuclease-free water. The purified DNA was amplified by 8 cycles of PCR using custom barcode primers (*76*) (**table S1**), purified again with the MinElute Reaction Cleanup Kit, and subjected to paired-end (2 × 150 bp) Illumina sequencing on a NovaSeq X Plus platform.

For RNA-seq, 2 × 10^6^ cells from each line were harvested before and 24 h after doxycycline treatment (1 µg/mL) as described above. After two washes with ice-cold PBS, total RNA was isolated using a Monarch Total RNA Miniprep Kit (NEB) according to the manufacturer’s protocol. mRNA was isolated from an average of 1 µg of total RNA by poly(dT) enrichment using the NEBNext Poly(A) mRNA Magnetic Isolation Module (NEB) according to the manufacturer’s instructions. cDNA was synthesized using NEBNext First Strand and Second Strand Synthesis Modules (NEB) according to the manufacturer’s instructions. Libraries were prepared using the NEBNext Ultra II DNA Library Prep Kit (NEB) following the manufacturer’s protocol. After purification with AMPure XP beads at a beads-to-sample ratio of 0.9:1 (v/v), libraries were quantified using a Fragment Analyzer (Agilent) and subjected to paired-end Illumina sequencing on a NovaSeq X Plus platform.

### Sequencing Data Processing and Analysis

Sequencing reads from ChIP-seq and MNase-ChIP-seq experiments were filtered using fastp v0.23.2 (*77*) with the following parameters: -g -q 20 -u 50 -n 5 -l 20 --overlap_diff_limit 1 -- overlap_diff_percent_limit 10. Reads were aligned to the human reference genome GRCh38 using Bowtie2 v2.3.4.1 (*78*) with the parameters: --end-to-end --very-sensitive --no-mixed --no-discordant --phred33 -I 10 -X 700. Unmapped reads and PCR duplicates were removed using samtools v1.9 (*79*) and Picard v2.26.11 (https://github.com/broadinstitute/picard). Peaks were called using the MACS2 *callpeak* command (v2.2.7.1) (*80*) with parameters -g hs -f BAMPE -- keep-dup all, using IgG or input samples as controls. Peaks from biological duplicates were assessed using IDR v2.0.3 (*81*), and concordant peaks were retained for downstream analysis*. De novo* motifs within ChIP-seq peaks were identified using the HOMER *findMotifsGenome.pl* command (v4.11.1) (*82*) with parameters -preparse -mask -size 50.

BAM files were normalized to 1× GRCh38 genome coverage (RPGC; -- effectiveGenomeSize 2913022398) using the bamCoverage tool from deepTools2 (*83*) with parameters --extendReads --binSize 10. Coverage depths were used to generate read-density heatmaps using the deepTools2 computeMatrix tool in reference-point mode with parameters -a 1000 -b 1000 -referencePoint center -missingDataAsZero. Sorted ChIP-seq or ATAC-seq peak BED files were used as reference regions (-R), and normalized ChIP-seq, ATAC-seq, and motif-site-derived bigWig files were used as signal tracks (-S). Heatmaps were generated from the resulting matrices using the deepTools2 plotHeatmap tool.

To plot the distribution of MNase-ChIP-seq fragment centers around CEBPB motif sites, fragments of 130–170 bp in length were used. Genomic sites recognized by CEBPB motifs were identified using MOODS v1.9.4.1 (*28*) with a P-value threshold of 0.0001 and an absolute score > 5. For each motif match overlapping ChIP-seq peaks, fragment centers were aggregated into consecutive 5-bp bins, and the total number of fragments in each bin was counted. Counts were normalized to the maximum value of the bin with the highest count, yielding values between 0 and 1. Data from all motif matches were aggregated, and the resulting profile was smoothed using kernel smoothing with a Gaussian kernel (bandwidth = 10).

RNA-seq reads were filtered using fastp v0.23.2 with parameters -g -q 20 -u 50 -n 10 -l 50 --detect_adapter_for_pe. Reads were then aligned to the GRCh38 reference genome using HISAT2 v2.1.0 (*84*). Gene-level counts for paired-end reads were quantified using featureCounts v1.6.0 (*85*) with the Ensembl gene annotation (release 86, GRCh38). Raw counts were normalized to FPKM (fragments per kilobase of transcript per million mapped fragments) to quantify gene expression levels. Differential expression analysis was performed using the R package DESeq2 (*86*). Differentially expressed genes were defined as those with FDR < 0.05 and |log (fold change)| ≥ 1. Genes with mean FPKM > 1 across biological replicates in at least one condition were selected for heatmap visualization and subsequent Gene Ontology (GO) enrichment analysis. GO enrichment was performed using the enrichGO function from the clusterProfiler R package (*87*) with parameters OrgDb = org.Hs.eg.db, keyType = “ENSEMBL”, ont = “ALL”, pAdjustMethod = “fdr”, and qvalueCutoff = 0.05.

### Drosophila Transgenic and Knock-in Strain Generation, Lifespan Analysis, and Genetic Ablation Assays

To generate transgenic and knock-in flies expressing Ets21C variants with point mutations (D287K, D289K, and N303W), we constructed three donor plasmids: Ets21C-WT-GFP and Ets21C-Mut1-GFP (for transgenic flies), and Ets21C-Mut2-GFP (for knock-in flies; see **table S8**). All constructs were generated by Gibson assembly and cloned into the *EcoRI* site of the pCaSpeR3 vector. Plasmids expressing *Ets21C*-targeting sgRNAs were synthesized and cloned into the pEASY-Blunt simple U6 vector (**table S8**) and subsequently used for knock-in fly generation. All constructs were verified by colony PCR and Sanger sequencing. Plasmids were submitted to the Core Facility of *Drosophila* Resource and Technology (Center for Excellence in Molecular Cell Science, Chinese Academy of Sciences) for embryo injection to generate transgenic and knock-in flies.

The *Drosophila melanogaster* w¹¹¹□ strain was used for knock-in experiments, yielding the w¹¹¹; Ets21C^KKW^-GFP line. For transgenic fly generation, the *Ets21C* knockout strain w¹¹¹; Ets21C^Δ10^ (a gift from the laboratory of Mirka Uhlirova, University of Cologne) was used. In these flies, either wild-type (w¹¹¹□; Ets21C^Δ10^; Ets21C^WT-GFP^) or mutant (w¹¹¹□; Ets21C^Δ10^; Ets21C^KKW^-GFP) *Ets21C* transgenes were site-specifically integrated at the attP2 landing site.

Unless otherwise stated, all flies were maintained at 25 °C on standard cornmeal-agar medium. For lifespan analysis, newly eclosed wild-type, knock-in, and transgenic flies (less than 1 day old) were collected (15 males and 15 females per vial) and transferred to fresh food daily. At 3 days of age, flies were transferred to vials containing a cotton plug soaked with 500 µL of either 5% sucrose (mock treatment) or 50 mM paraquat in 5% sucrose (PQ). Mortality was scored every 6 hours. The assay was performed with four biological replicates (**fig. S9** and **table S8**). Survival curves were generated from the mortality data and compared using the log-rank test in R.

Genetic ablation experiments were performed as described by Worley *et al*. (*50*). Briefly, flies of the w¹¹¹; Ets21C^Δ10^ strain were crossed to rn-GAL4, tub-GAL80^ts^, UAS-egr transgenic flies (Bloomington Drosophila Stock Center, 51280) to generate w¹¹¹; Ets21C^Δ10^; rn-GAL4, tub-GAL80^ts^, UAS-egr flies. Virgin females of the w¹¹¹; Ets21C^Δ10^; rn-GAL4, tub-GAL80^ts^, UAS-egr genotype were then crossed to transgenic males carrying either wild-type (w¹¹¹; Ets21C^Δ10^; Ets21C^WT^-GFP) or mutant (w¹¹¹; Ets21C^Δ10^; Ets21C^KKW^-GFP) *Ets21C* transgenes. This produced progeny of genotypes w¹¹¹; Ets21C^Δ10^; rn-GAL4, tub-GAL80^ts^, UAS-egr / Ets21C^WT^-GFP and w¹¹¹; Ets21C^Δ10^; rn-GAL4, tub-GAL80^ts^, UAS-egr / Ets21C^KKW^-GFP, which were used for genetic ablation assays. 0–24-hour embryos were collected and maintained at 25 °C for approximately 3.5 days until they reached the wandering third-instar larval stage.

Synchronized larvae were then shifted from 25 °C to 30 °C and maintained at 30 °C for 40 hours to induce genetic ablation in wing imaginal discs. Pupae were then returned to 25 °C to allow tissue regeneration. Adult wings were examined under a stereomicroscope. Regeneration efficiency was scored by assigning each wing to one of five categories based on the percentage of tissue recovered (*88*): 0% − 10%, 10% − 30%, 30% − 50%, 50% − 70%, or ≥70% (**fig. S9** and **table S8**). The experiment was performed with three biological replicates. For imaging, adult flies were anesthetized with CO, transferred to empty vials to minimize ice crystal formation, and frozen at −80 °C for 30 min. Wing images were acquired using a Sunny Optical DMS1000 3D digital microscope.

**Fig. S1.**
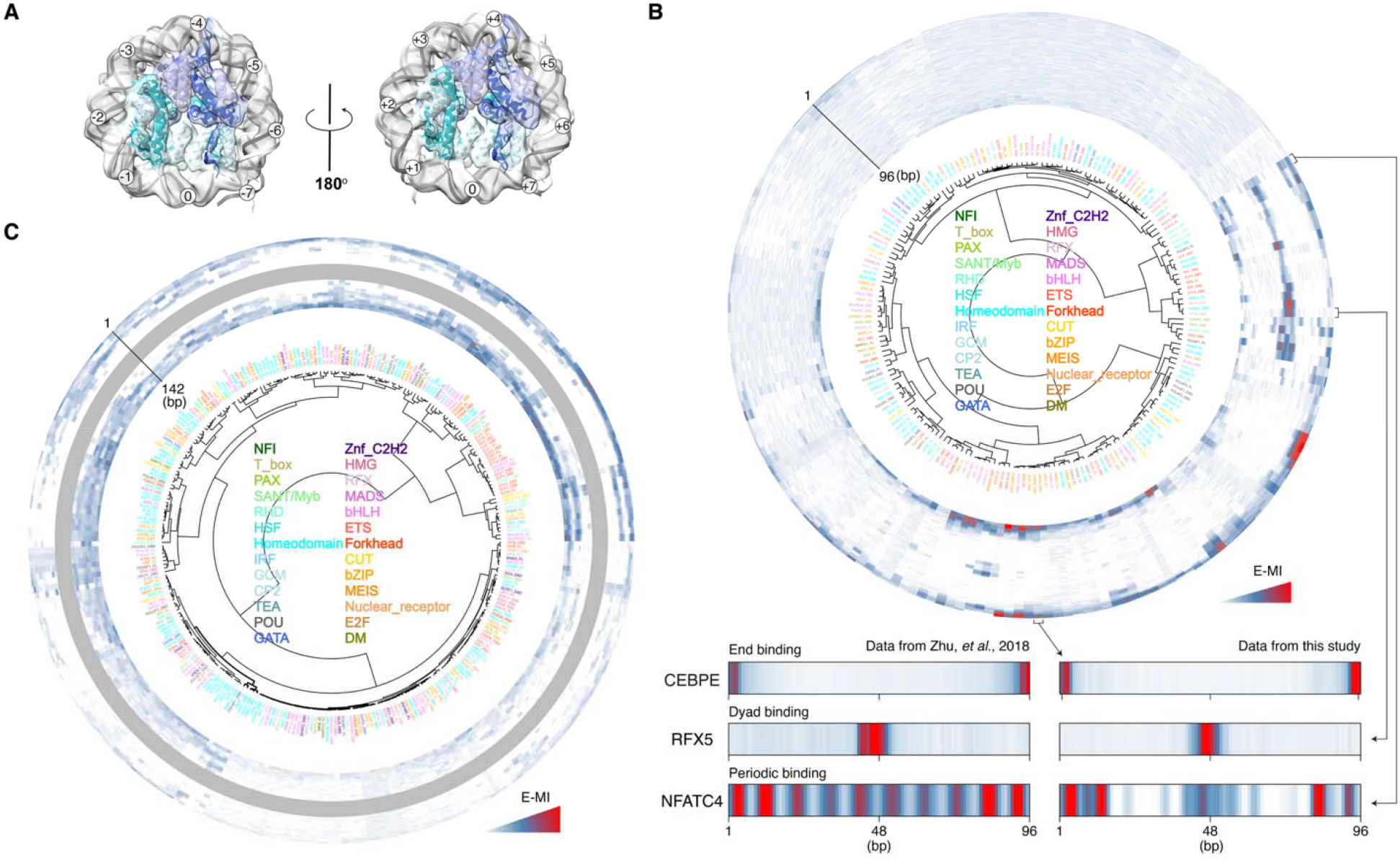
Positional preferences of TFs on nucleosomal DNA. (**A**) Cryo-EM map and corresponding model of the nucleosome with the 193-bp ligand (lig193). Two top views are shown, related by a 180° rotation. (**B** and **C**) Hierarchical clustering of E-MI diagonals for NCAP-SELEX with the 147-bp ligand (lig147; **B**) and 193-bp ligand (lig193; **C**). TFs are color-coded by family. Examples of TFs showing end-, periodic-, and dyad-preference binding modes are indicated (**table S3**). Similar preferences were also observed in our previous study (*26*).

**Fig. S2.**
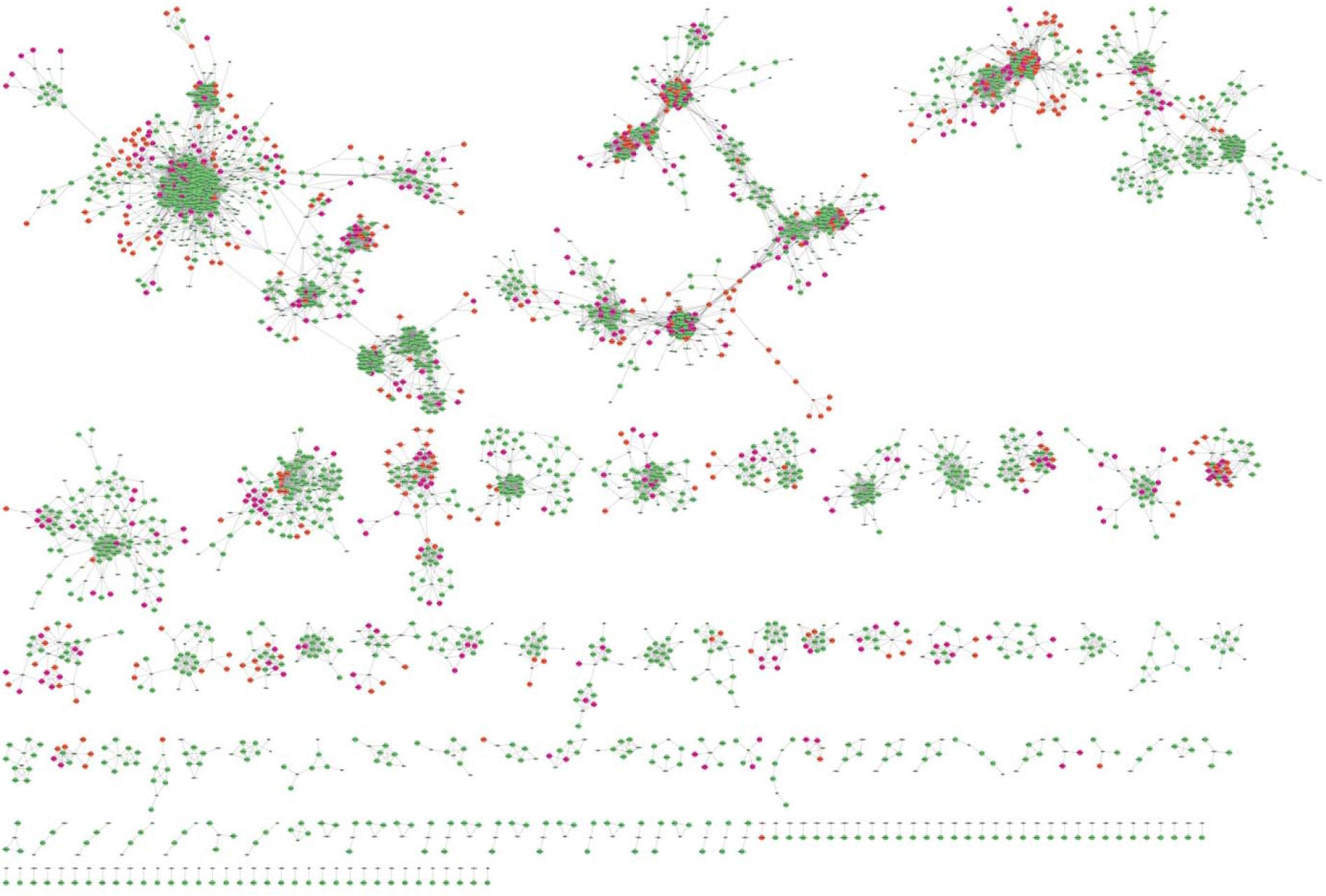
Motif similarity analysis. Network analysis of the similarity of NCAP-SELEX motifs (red) to HT-SELEX data (green). Edges are drawn between the motifs if they have Ssum similarity > 1.5 × 10^−5^ (SSTAT (*57*)) and the similarities are visualized using Cytoscape (version v3.10.1(*58*)).

**Fig. S3.**
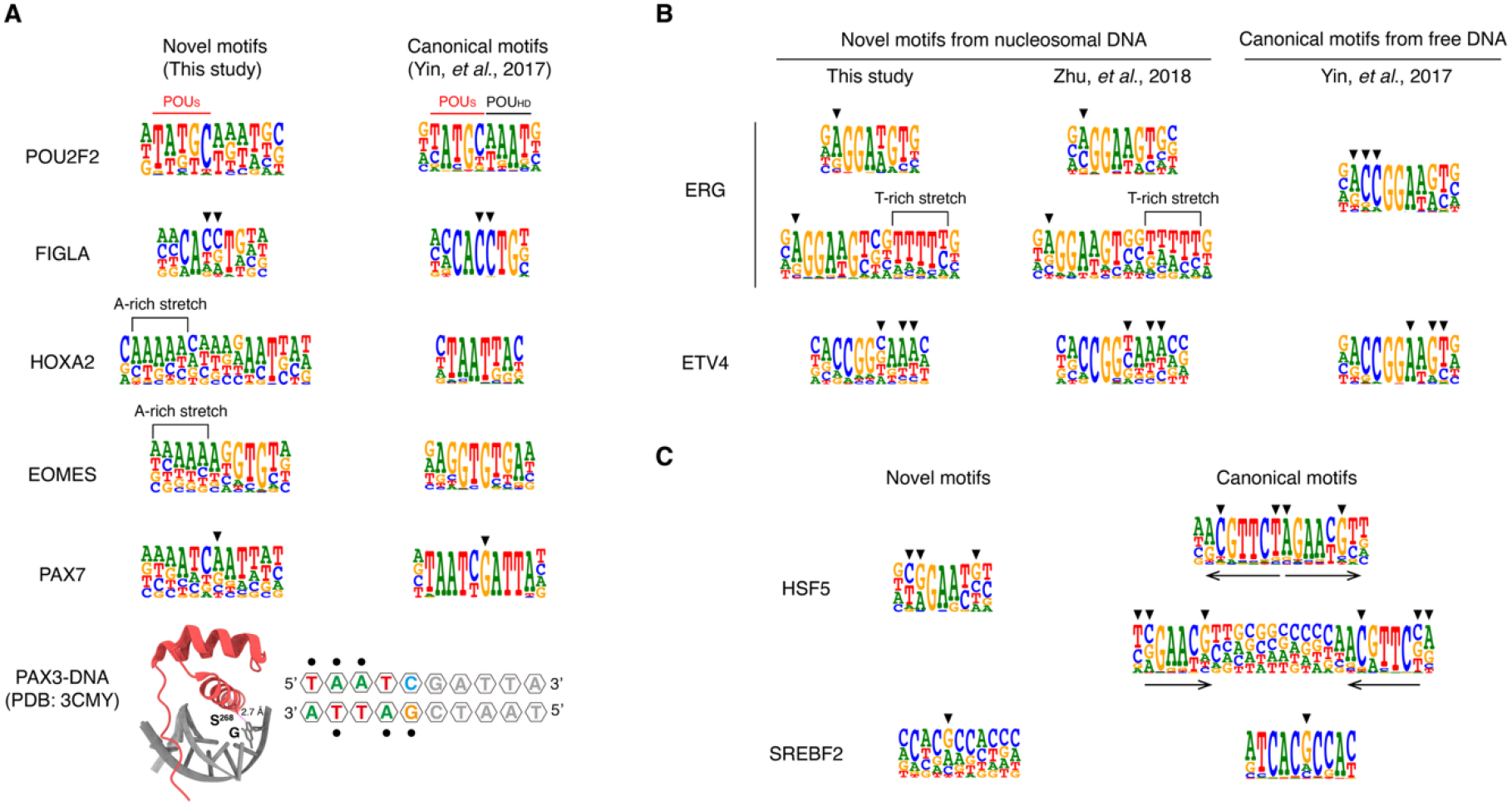
Representative novel motifs identified by NCAP-SELEX from nucleosomal DNA. **(A**-**C)** Motif logos for TFs whose NCAP-SELEX motifs differ from their corresponding HT-SELEX counterparts. Left panels in **A** and **C** show novel motifs recovered from nucleosomal DNA (NCAP-SELEX); right panels show canonical motifs recovered from free DNA (HT-SELEX). The structure in **A** shows the interaction between PAX3 and free DNA (PDB entry 3CMY); dots indicate bases that contact the TF protein through hydrogen bonds. In **B**, NCAP-SELEX motifs derived in this study (left) are compared with motifs identified in previous studies (*25, 26*) using NCAP-SELEX (middle) or HT-SELEX (right). Arrowheads indicate positions where TFs recognize distinct bases on nucleosomal versus free DNA.

**Fig. S4.**
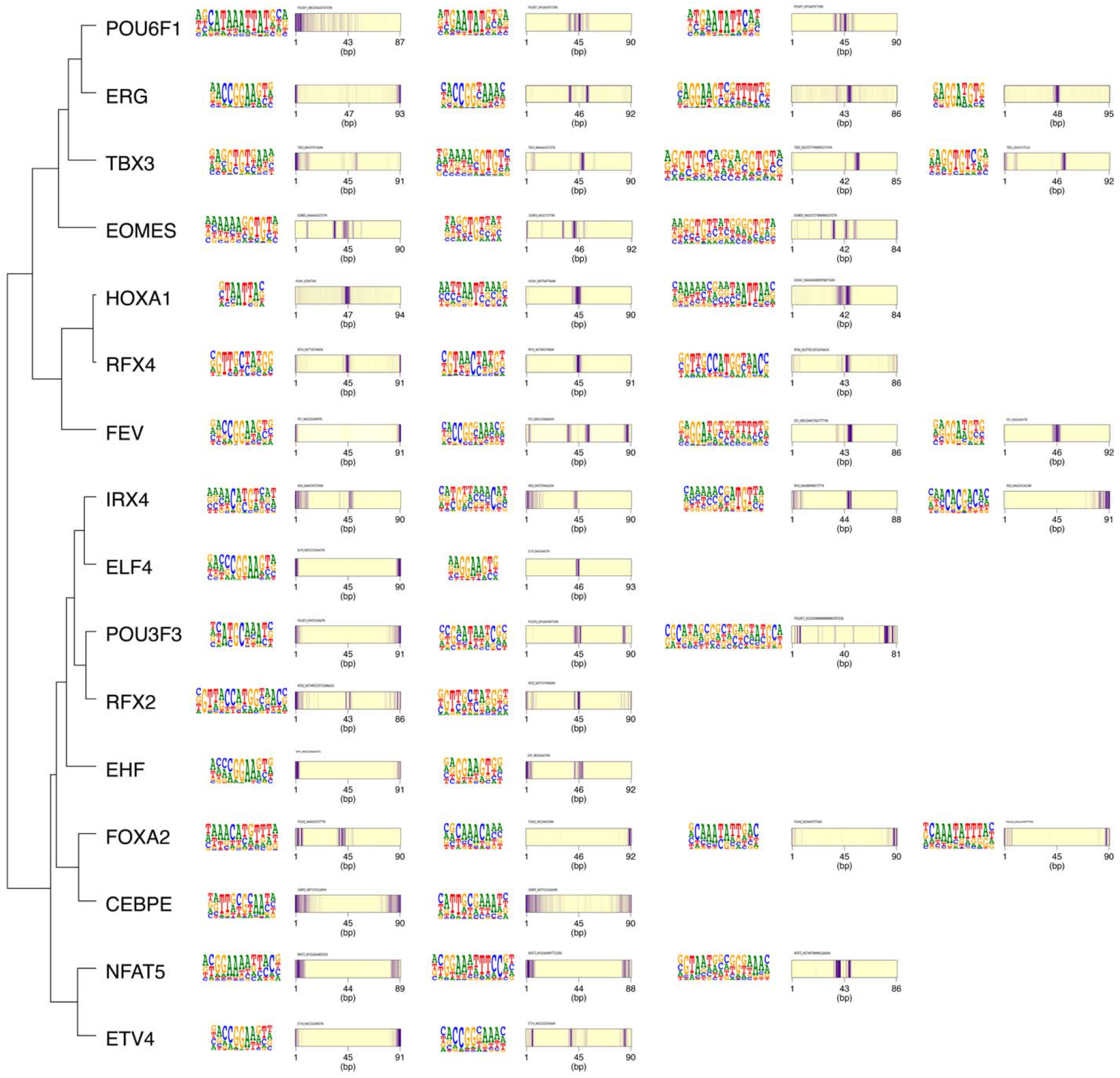
TFs exhibiting diverse binding modes and specificities. Motif logos of NCAP-SELEX PWMs and their motif-matching results on lig147 are shown. Hierarchical clustering was performed as in Fig. 2H, based on E-MI diagonals of individual TFs. Underlying data are provided in **table S3**.

**Fig. S5.**
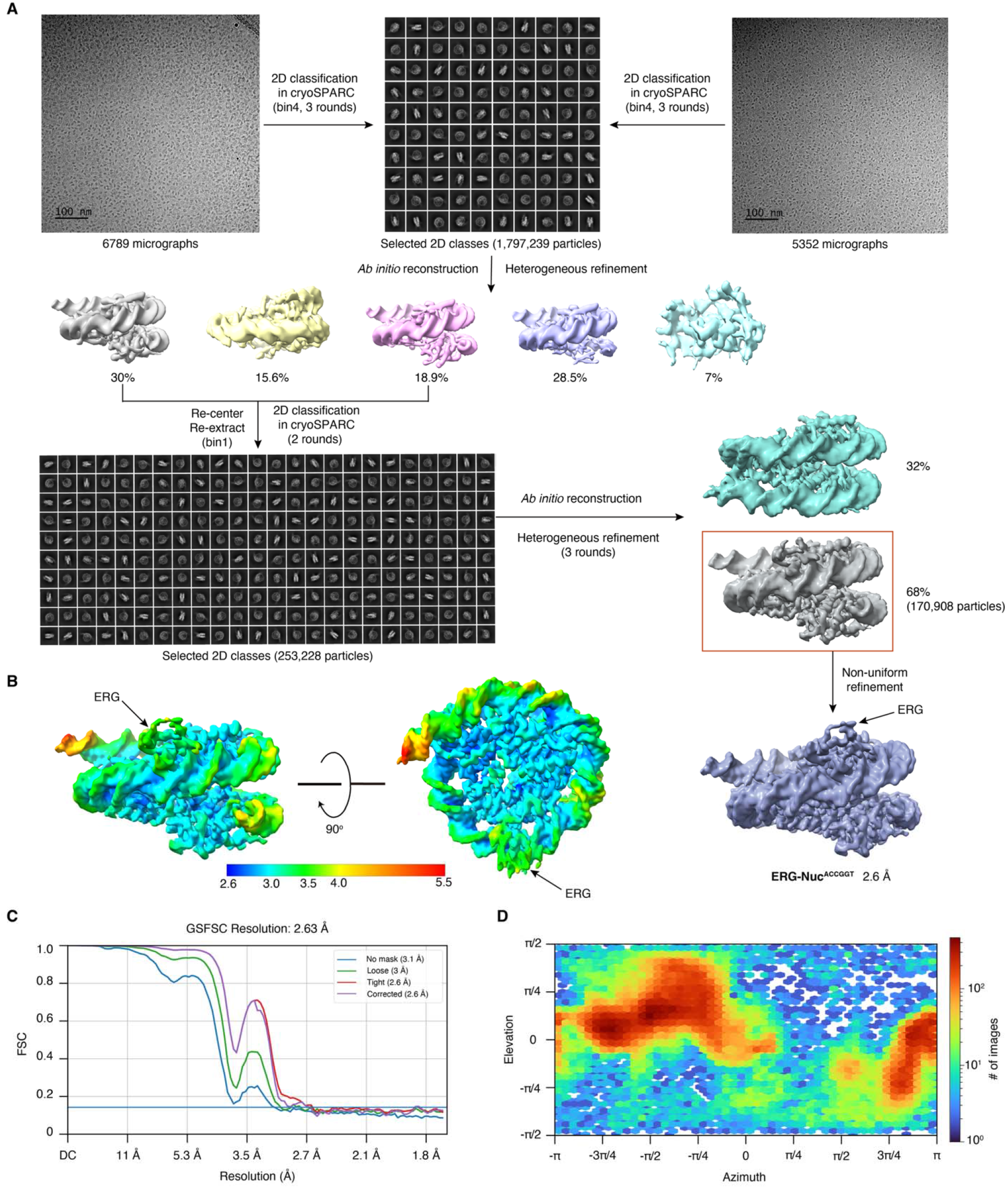
Cryo-EM processing of the ERG–Nuc^ACCGGT^ complex. (**A**) Schematic workflow for 3D reconstruction of the ERG–Nuc^ACCGGT^ complex. Representative raw cryo-EM micrographs are shown in the upper left and upper right panels. A total of 12,141 micrographs were collected using a Krios G4 300 keV microscope in two batches (6,789 and 5,353 micrographs, respectively). All dose-fractionated stacks were subjected to beam-induced motion correction using MotionCor2 within RELION 3.1 and then transferred to cryoSPARC v5.0.6 for subsequent processing. Particles were picked using the Blob picker, followed by three rounds of 2D classification to remove poor-quality particles prior to ab initio reconstruction. After one round of heterogeneous refinement of the five ab initio classes, two classes displaying complete nucleosome features were retained; the remaining three classes were discarded. Particles from the two good classes were re-extracted without binning and subjected to two additional rounds of 2D classification. High-resolution classes were selected for a new round of ab initio reconstruction. After three rounds of heterogeneous refinement, two models were obtained, one of which showed density corresponding to ERG, accounting for 68% of the particles. Refinement of the best particles using non-uniform refinement in cryoSPARC yielded a map at 2.6 Å resolution. (**B**) Local resolution map of the ERG–Nuc^ACCGGT^ complex. The map is color-coded by local resolution, ranging from 2.6 Å (blue) to 5.5 Å (red). (**C**) Fourier shell correlation (FSC) curves of the maps, with the FSC = 0.143 cutoff indicated. (**D**) Angular distribution of particles used for reconstruction.

**Fig. S6.**
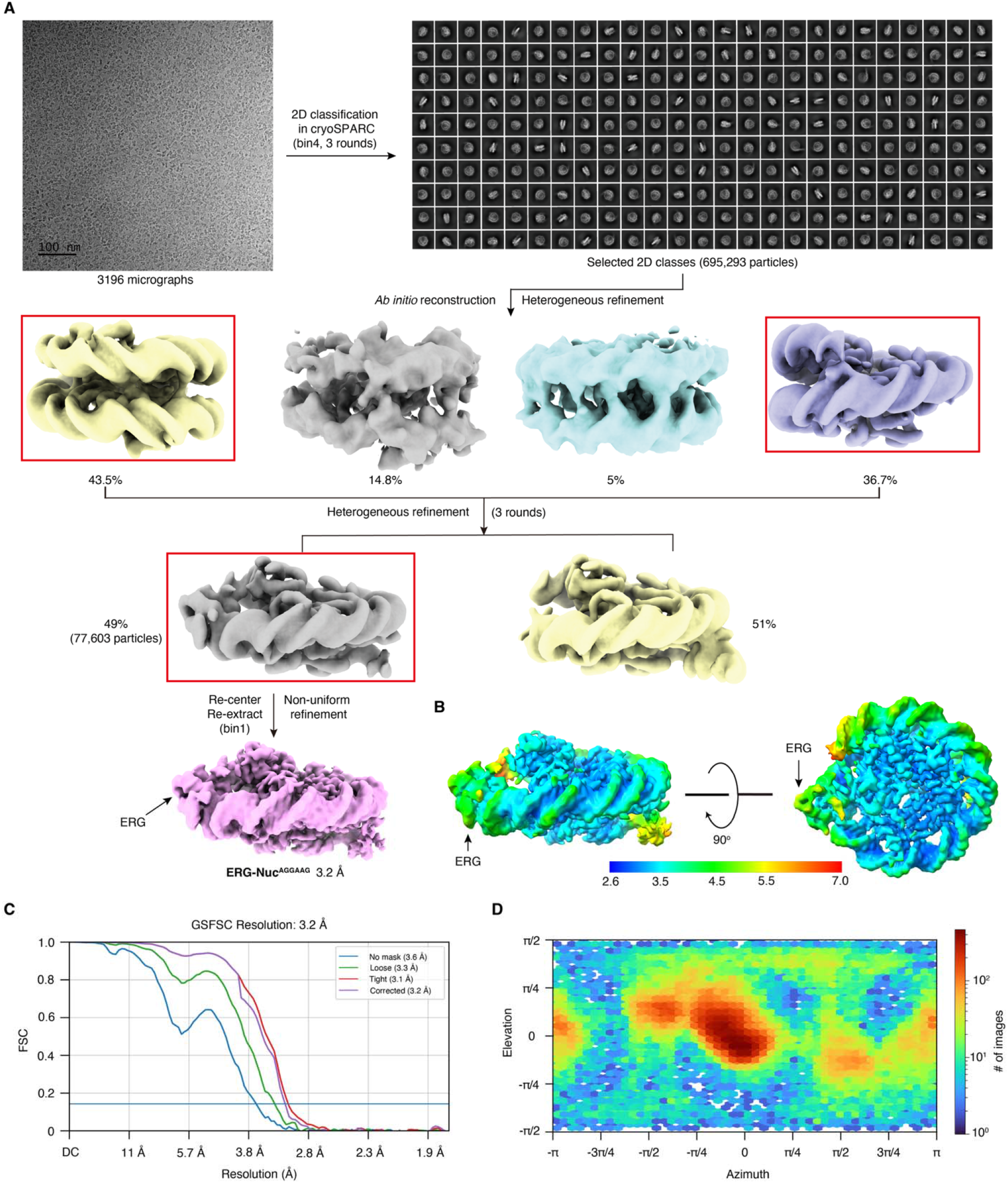
Cryo-EM processing of the ERG–Nuc^AGGAAG^ complex. (**A**) Schematic workflow for 3D reconstruction of the ERG–Nuc^AGGAAG^ complex. The upper left panel shows a representative cryo-EM micrograph from a set of 3,196 micrographs collected on a Titan Krios microscope at 300 keV; the upper right panel shows representative 2D class averages. All dose-fractionated stacks were subjected to beam-induced motion correction using MotionCor2 within RELION 3.1 and then transferred to cryoSPARC v5.0.6 for subsequent processing. Particles were picked using the Blob picker, followed by three rounds of 2D classification to remove poor-quality particles prior to ab initio reconstruction. After one round of heterogeneous refinement of the four ab initio classes, two classes with complete nucleosome features were retained; the remaining classes were discarded. Particles from the two good classes were re-extracted without binning. After three rounds of heterogeneous refinement, two models were obtained, one of which showed density corresponding to ERG, accounting for 49% of the particles. Refinement of the best particles using non-uniform refinement in cryoSPARC yielded a map at 3.2 Å resolution. (**B**) Local resolution map of the ERG–Nuc^AGGAAG^ complex. The map is color-coded by local resolution, ranging from 2.6 Å (blue) to 7.0 Å (red). (**C**) Fourier shell correlation (FSC) curves of the maps, with the FSC = 0.143 cutoff indicated. (**D**) Angular distribution of particles used for reconstruction.

**Fig. S7.**
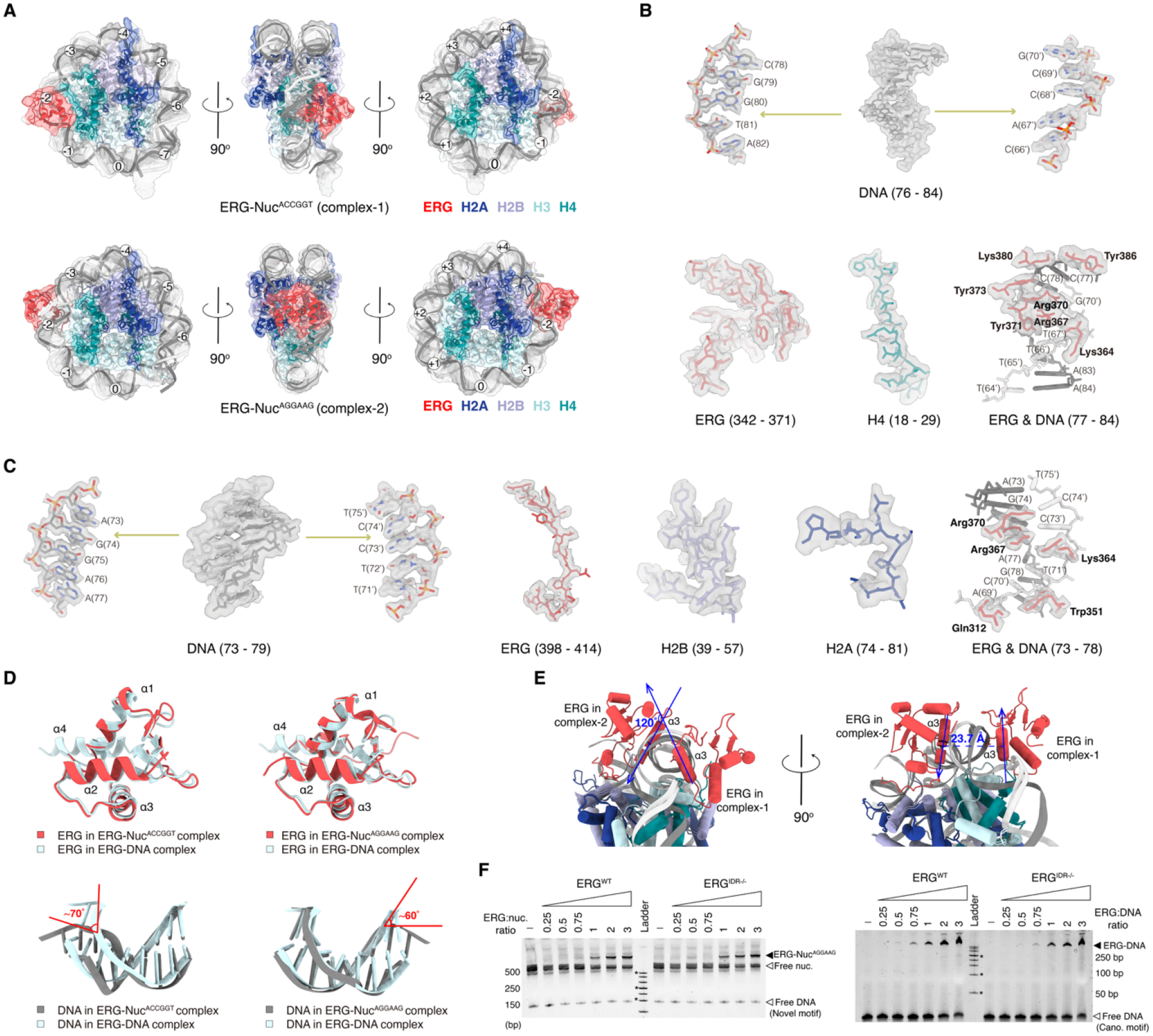
Cryo-EM model details of the ERG–Nuc^ACCGGT^ and ERG–Nuc^AGGAAG^ complexes. (**A**) Cryo-EM maps and corresponding atomic models of the ERG–Nuc^ACCGGT^ (upper) and ERG–Nuc^AGGAAG^ (lower) complexes. SHL positions are labeled. Cryo-EM maps are colored as in Fig. 3 and shown at 50% transparency. For each complex, three views related by 90° rotations around the dyad are shown (left, middle, and right). (**B**) Cryo-EM maps and corresponding models of the ERG–Nuc^ACCGGT^ complex, showing representative regions: the ERG-binding motif, the ERG–H4 interaction interface, and ERG residues that contact nucleosomal DNA. (**C**) Cryo-EM maps and corresponding models of the ERG–Nuc^AGGAAG^ complex, showing representative regions: the ERG-binding motif, the ERG–H2A/H2B interaction interface, and ERG residues that contact nucleosomal DNA. (**D**) Superposition of the ERG–Nuc complexes with the ERG–DNA complex (PDB entry 5YBD). Upper panels: superposition of ERG from ERG–Nuc^ACCGGT^ (left) and ERG–Nuc^AGGAAG^ (right) with ERG from the ERG–DNA complex. Lower panels: superposition of DNA (binding motif) from ERG–Nuc^ACCGGT^ (left) and ERG–Nuc^AGGAAG^ (right) with DNA from the ERG–DNA complex. (**E**) Superposition of the ERG–Nuc^ACCGGT^ and ERG–Nuc^AGGAAG^ models. Left: superposition of ERG structures showing that the α3 helices of the two ERG molecules form a 120° angle. Right: superposition showing the horizontal distance between the α3 helix centers is 23.7 Å. (**F**) Representative EMSA showing binding of increasing amounts of wild-type (ERG^WT^) and mutant (ERG^IDR−/−^, deletion of the intrinsically disordered region (IDR; residues 399–414)) ERG to nucleosomes (left) and free DNA (right). Black arrowheads indicate ERG–nucleosome (left) and ERG–DNA (right) complexes; white arrowheads indicate free DNA and free nucleosomes. The DNA in the nucleosome binding assay (left) contains the novel motif consensus (AGGAAGT), whereas the DNA in the DNA binding assay (right) contains the canonical motif consensus (ACCGGAAGT).

**Fig. S8.**
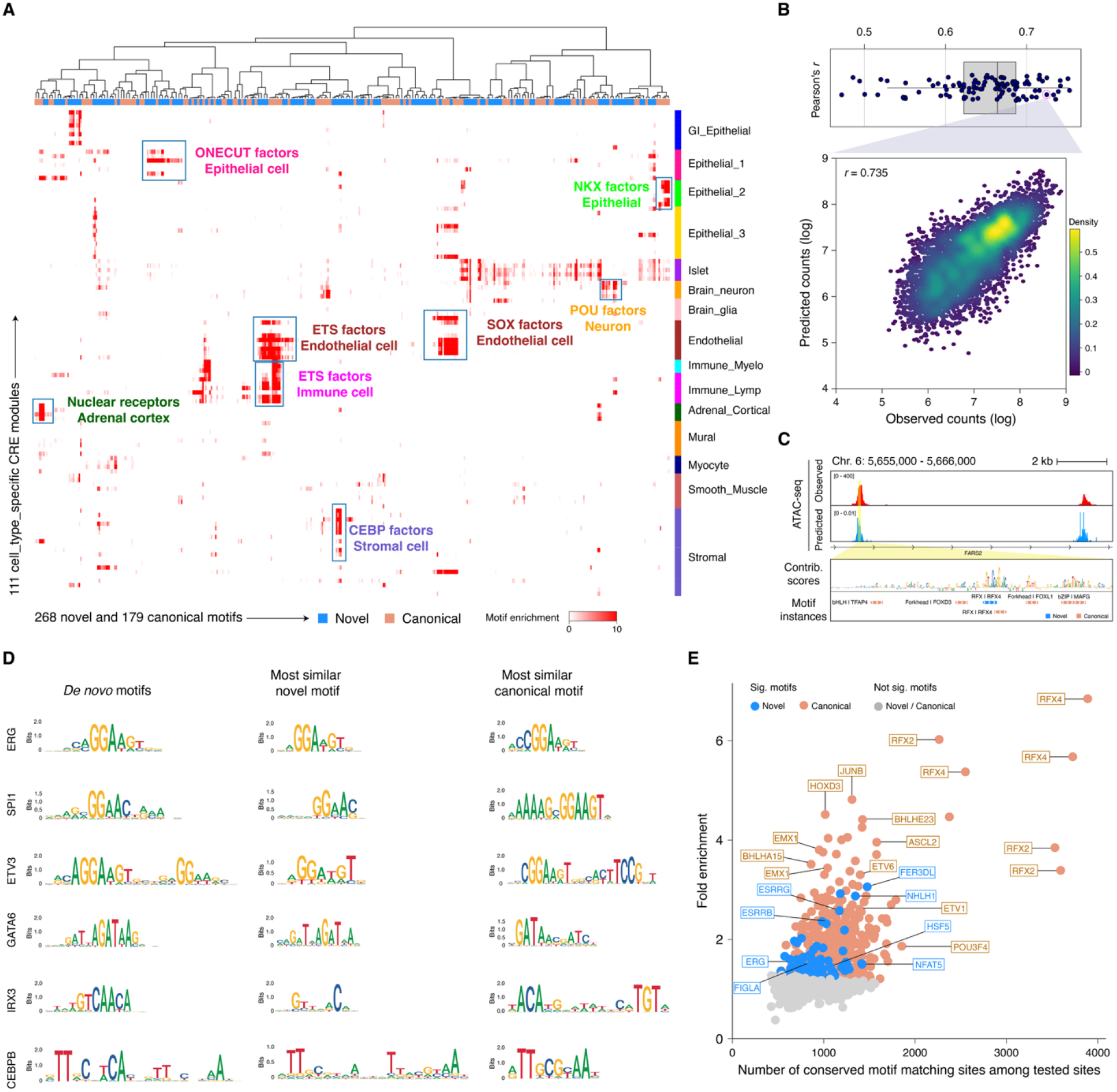
Comprehensive analysis of novel motif properties. (**A**) Hierarchically clustered heatmap showing enrichment (−log_10_(q values); see **Methods**) of novel and corresponding canonical motifs in cell-type-specific cCREs from CATlas (*34*). Rows represent samples; columns represent motifs. (**B**) Top: distribution of Pearson’s r between predicted and observed log-transformed counts in the selected 30,000 peak regions per model (averaged across five folds). Bottom: predicted (y-axis) versus observed (x-axis) log-transformed counts for one model fold from endothelial cells. (**C**) Example of the FARS2 locus in alpha-1 islet cells, showing observed and predicted chromatin accessibility, per-base contribution scores, and annotated motif instances. (**D**) Examples of *de novo* motifs derived from the CATlas dataset (*34*) using ChromBPNet that are more similar to NCAP-SELEX than to HT-SELEX motifs. Left to right: *de novo* motif representation as a contribution weight matrix (CWM), most similar NCAP-SELEX motif (PWM), and most similar HT-SELEX motif (PWM). (**E**) Motif conservation analysis. The y-axis shows fold enrichment, calculated as the fraction of conserved matches for the indicated motif divided by the fraction of conserved matches for its control motif (see **Methods**). The x-axis shows the number of conserved motif matches among the selected sites. For each motif, 10,000 non-overlapping, highest-affinity sites within Zoonomia-defined human constrained non-coding regions were selected for both the true motif and its control motifs. Conservation significance was determined using a hypergeometric test, with P-values adjusted to control the family-wise error rate (FWER). Motifs with significant conservation (adjusted P < 0.05) are colored in blue (novel NCAP-SELEX motifs) or orange (canonical HT-SELEX motifs); motifs without significant conservation are shown in grey. Selected motifs with the lowest P-values in each category are labeled with TF symbols. Underlying data, including corrected P-values, are provided in **table S9**.

**Fig. S9.**
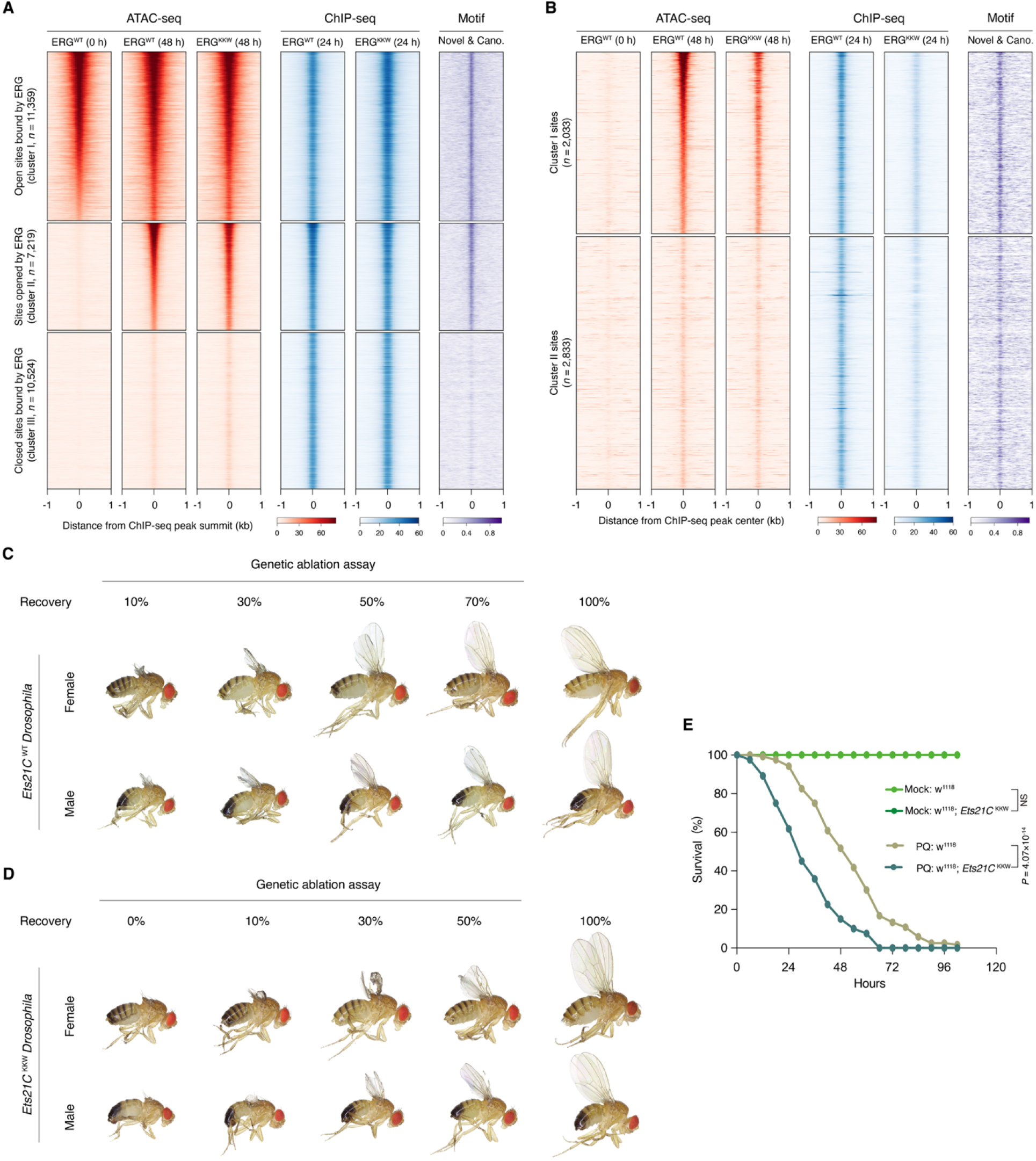
Biological roles of ERG nucleosome-binding capacity. (**A**) Read-density heatmaps showing normalized ATAC-seq signal (red), ChIP-seq intensity (blue) and motif enrichment (purple) for ERG^WT^ and ERG^KKW^ in H9 human ES cells. ATAC-seq signals before and 48 h after ectopic induction of ERG^WT^ and ERG^KKW^ are shown. Chromatin sites are ranked by ATAC-seq signal and divided into three clusters: cluster I, sites that are open and bound by ERG; cluster II, sites that become open after ERG expression; and cluster III, sites that are bound by ERG but remain closed 48 h after ERG expression. Cluster sizes and color scales are indicated at the bottom. (**B**) As in **A**, but for ERG^WT^-specific ChIP-seq peaks. Sites are divided into two clusters: sites that become open (top, cluster I) or remain closed (bottom, cluster II) 48 h after ERG^WT^ induction. (**C** and **D**) Adult wing sizes in *Ets21C*^WT^ (**C**) and *Ets21C*^KKW^ (**D**) *Drosophila* after the ablation protocol, compared with normal wings. Left: extent of adult wing regeneration, scored by binning wings into five categories (0% - 10%, 10% - 30%, 30% - 50%, 50% - 70%, or ³70%). (**E**) As in Fig. 5F, but for knock-in flies. Survival curves were derived from four independent experiments. Statistical significance was determined by the log-rank test. NS, not significant.

## References and Notes

1. S. A. Lambert et al., The Human Transcription Factors. Cell 175, 598–599 (2018).

2. Z. Wunderlich, L. A. Mirny, Different gene regulation strategies revealed by analysis of binding motifs. Trends Genet 25, 434–440 (2009).

3. S. Kim, J. Wysocka, Deciphering the multi-scale, quantitative cis-regulatory code. Mol Cell 83, 373–392 (2023).

4. C. G. de Boer, J. Taipale, Hold out the genome: a roadmap to solving the cis-regulatory code. Nature 625, 41–50 (2024).

5. Z. Xie et al., DNA-guided transcription factor interactions extend human gene regulatory code. Nature 641, 1329–1338 (2025).

6. P. Blomquist, Q. Li, O. Wrange, The affinity of nuclear factor 1 for its DNA site is drastically reduced by nucleosome organization irrespective of its rotational or translational position. Journal of Biological Chemistry 271, 153–159 (1996).

7. K. Luger, A. W. Mader, R. K. Richmond, D. F. Sargent, T. J. Richmond, Crystal structure of the nucleosome core particle at 2.8 A resolution. Nature 389, 251–260 (1997).

8. A. K. Michael, N. H. Thoma, Reading the chromatinized genome. Cell 184, 3599–3611 (2021).

9. Q. Li, O. Wrange, Translational positioning of a nucleosomal glucocorticoid response element modulates glucocorticoid receptor affinity. Genes Dev 7, 2471–2482 (1993).

10. A. Soufi et al., Pioneer transcription factors target partial DNA motifs on nucleosomes to initiate reprogramming. Cell 161, 555–568 (2015).

11. K. S. Zaret, S. E. Mango, Pioneer transcription factors, chromatin dynamics, and cell fate control. Curr Opin Genet Dev 37, 76–81 (2016).

12. A. Mayran et al., Pioneer factor Pax7 deploys a stable enhancer repertoire for specification of cell fate. Nat Genet 50, 259–269 (2018).

13. M. L. Bulyk, J. Drouin, M. M. Harrison, J. Taipale, K. S. Zaret, Pioneer factors - key regulators of chromatin and gene expression. Nature Reviews Genetics 24, 809–815 (2023).

14. M. Carminati, L. Vecchia, L. Stoos, N. H. Thomä, Pioneer factors: Emerging rules of engagement for transcription factors on chromatinized DNA. Curr Opin Struc Biol 88, (2024).

15. T. J. Richmond, C. A. Davey, The structure of DNA in the nucleosome core. Nature 423, 145–150 (2003).

16. S. O. Dodonova, F. Zhu, C. Dienemann, J. Taipale, P. Cramer, Nucleosome-bound SOX2 and SOX11 structures elucidate pioneer factor function. Nature 580, 669–672 (2020).

17. A. K. Michael et al., Mechanisms of OCT4-SOX2 motif readout on nucleosomes. Science 368, 1460–1465 (2020).

18. A. K. Michael et al., Cooperation between bHLH transcription factors and histones for DNA access. Nature 619, 385-+ (2023).

19. M. Iwafuchi et al., Gene network transitions in embryos depend upon interactions between a pioneer transcription factor and core histones. Nat Genet 52, 418–427 (2020).

20. M. P. Meers, D. H. Janssens, S. Henikoff, Pioneer Factor-Nucleosome Binding Events during Differentiation Are Motif Encoded. Molecular Cell 75, 562-+ (2019).

21. H. Tanaka et al., Interaction of the pioneer transcription factor GATA3 with nucleosomes. Nat Commun 11, 4136 (2020).

22. R. Guan, T. Lian, B. R. Zhou, D. Wheeler, Y. Bai, Structural mechanism of LIN28B nucleosome targeting by OCT4. Mol Cell 83, 1970–1982 e1976 (2023).

23. A. Jolma et al., DNA-Binding Specificities of Human Transcription Factors. Cell 152, 327–339 (2013).

24. H. S. Najafabadi et al., C2H2 zinc finger proteins greatly expand the human regulatory lexicon. Nat Biotechnol 33, 555–562 (2015).

25. Y. Yin et al., Impact of cytosine methylation on DNA binding specificities of human transcription factors. Science 356, (2017).

26. F. Zhu et al., The interaction landscape between transcription factors and the nucleosome. Nature 562, 76–81 (2018).

27. W. Xue et al., Structural basis of nucleosome binding and destabilization by the extended DNA binding domain of RFX5. Nucleic Acids Res 53, (2025).

28. J. Korhonen, P. Martinmaki, C. Pizzi, P. Rastas, E. Ukkonen, MOODS: fast search for position weight matrix matches in DNA sequences. Bioinformatics 25, 3181–3182 (2009).

29. K. R. Nitta et al., Conservation of transcription factor binding specificities across 600 million years of bilateria evolution. Elife 4, (2015).

30. M. Nishimura, Y. Takizawa, K. Nozawa, H. Kurumizaka, Structural basis for p53 binding to its nucleosomal target DNA sequence. PNAS Nexus 1, pgac177 (2022).

31. I. Rauluseviciute et al., JASPAR 2024: 20th anniversary of the open-access database of transcription factor binding profiles. Nucleic Acids Res 52, D174–D182 (2024).

32. R. Sharma, S. P. Gangwar, A. K. Saxena, Comparative structure analysis of the ETSi domain of ERG3 and its complex with the E74 promoter DNA sequence. Corrigendum. Acta Crystallogr F Struct Biol Commun 75, 397–398 (2019).

33. C. D. Cooper, J. A. Newman, H. Aitkenhead, C. K. Allerston, O. Gileadi, Structures of the Ets Protein DNA-binding Domains of Transcription Factors Etv1, Etv4, Etv5, and Fev: DETERMINANTS OF DNA BINDING AND REDOX REGULATION BY DISULFIDE BOND FORMATION. J Biol Chem 290, 13692–13709 (2015).

34. K. Zhang et al., A single-cell atlas of chromatin accessibility in the human genome. Cell 184, 5985–6001 e5919 (2021).

35. S. J. Loughran et al., The transcription factor Erg is essential for definitive hematopoiesis and the function of adult hematopoietic stem cells. Nat Immunol 9, 810–819 (2008).

36. V. Kalna et al., The Transcription Factor ERG Regulates Super-Enhancers Associated With an Endothelial-Specific Gene Expression Program. Circ Res 124, 1337–1349 (2019).

37. C. K. McClard et al., POU6f1 Mediates Neuropeptide-Dependent Plasticity in the Adult Brain. J Neurosci 38, 1443–1461 (2018).

38. A. Pampari et al., ChromBPNet: bias factorized, base-resolution deep learning models of chromatin accessibility reveal cis-regulatory sequence syntax, transcription factor footprints and regulatory variants. bioRxiv, (2025).

39. B. B. Liu et al., Multiomics and deep learning dissect regulatory syntax in human development. Nature, (2026).

40. A. Shrikumar, P. Greenside, A. Kundaje, Learning Important Features Through Propagating Activation Differences. Proceedings of the 34th International Conference on Machine Learning, (2017).

41. C. Cheng et al., Identification of Rfx6 target genes involved in pancreas development and insulin translation by ChIP-seq. Biochem Biophys Res Commun 508, 556–562 (2019).

42. B. Memon et al., RFX3 is essential for the generation of functional human pancreatic islets from stem cells. Diabetologia 68, 1476–1491 (2025).

43. J. R. Hesselberth et al., Global mapping of protein-DNA interactions in vivo by digital genomic footprinting. Nat Methods 6, 283–289 (2009).

44. J. Vierstra et al., Global reference mapping of human transcription factor footprints. Nature 583, 729–736 (2020).

45. P. Vijayaraj et al., Erg is a crucial regulator of endocardial-mesenchymal transformation during cardiac valve morphogenesis. Development 139, 3973–3985 (2012).

46. G. M. Birdsey et al., The endothelial transcription factor ERG promotes vascular stability and growth through Wnt/beta-catenin signaling. Dev Cell 32, 82–96 (2015).

47. M. Bosch, F. Serras, E. Martin-Blanco, J. Baguna, JNK signaling pathway required for wound healing in regenerating Drosophila wing imaginal discs. Dev Biol 280, 73–86 (2005).

48. C. Bergantinos, M. Corominas, F. Serras, Cell death-induced regeneration in wing imaginal discs requires JNK signalling. Development 137, 1169–1179 (2010).

49. J. Mundorf, C. D. Donohoe, C. D. McClure, T. D. Southall, M. Uhlirova, Ets21c Governs Tissue Renewal, Stress Tolerance, and Aging in the Drosophila Intestine. Cell Rep 27, 3019–3033 e3015 (2019).

50. M. I. Worley et al., Ets21C sustains a pro-regenerative transcriptional program in blastema cells of Drosophila imaginal discs. Curr Biol 32, 3350–3364 e3356 (2022).

51. R. Vincentelli et al., High-throughput protein expression screening and purification in Escherichia coli. Methods 55, 65–72 (2011).

52. K. Struhl, E. Segal, Determinants of nucleosome positioning. Nat Struct Mol Biol 20, 267–273 (2013).

53. Z. Gu, L. Gu, R. Eils, M. Schlesner, B. Brors, circlize Implements and enhances circular visualization in R. Bioinformatics 30, 2811–2812 (2014).

54. A. Jolma et al., Multiplexed massively parallel SELEX for characterization of human transcription factor binding specificities. Genome Res 20, 861–873 (2010).

55. A. Jolma et al., DNA-dependent formation of transcription factor pairs alters their binding specificity. Nature 527, 384–388 (2015).

56. S. Gupta, J. A. Stamatoyannopoulos, T. L. Bailey, W. S. Noble, Quantifying similarity between motifs. Genome Biol 8, R24 (2007).

57. U. J. Pape, S. Rahmann, M. Vingron, Natural similarity measures between position frequency matrices with an application to clustering. Bioinformatics 24, 350–357 (2008).

58. P. Shannon et al., Cytoscape: a software environment for integrated models of biomolecular interaction networks. Genome Res 13, 2498–2504 (2003).

59. M. J. Christmas et al., Evolutionary constraint and innovation across hundreds of placental mammals. Science 380, eabn3943 (2023).

60. P. F. Sullivan et al., Leveraging base-pair mammalian constraint to understand genetic variation and human disease. Science 380, eabn2937 (2023).

61. G. M. Cooper et al., Distribution and intensity of constraint in mammalian genomic sequence. Genome Res 15, 901–913 (2005).

62. K. S. Pollard, M. J. Hubisz, K. R. Rosenbloom, A. Siepel, Detection of nonneutral substitution rates on mammalian phylogenies. Genome Res 20, 110–121 (2010).

63. P. N. Dyer et al., Reconstitution of nucleosome core particles from recombinant histones and DNA. Methods Enzymol 375, 23–44 (2004).

64. S. Q. Zheng et al., MotionCor2: anisotropic correction of beam-induced motion for improved cryo-electron microscopy. Nat Methods 14, 331–332 (2017).

65. J. Zivanov et al., New tools for automated high-resolution cryo-EM structure determination in RELION-3. Elife 7, (2018).

66. A. Rohou, N. Grigorieff, CTFFIND4: Fast and accurate defocus estimation from electron micrographs. J Struct Biol 192, 216–221 (2015).

67. A. Punjani, J. L. Rubinstein, D. J. Fleet, M. A. Brubaker, cryoSPARC: algorithms for rapid unsupervised cryo-EM structure determination. Nat Methods 14, 290–296 (2017).

68. E. F. Pettersen et al., UCSF Chimera--a visualization system for exploratory research and analysis. J Comput Chem 25, 1605–1612 (2004).

69. P. Emsley, K. Cowtan, Coot: model-building tools for molecular graphics. Acta Crystallogr D Biol Crystallogr 60, 2126–2132 (2004).

70. P. D. Adams et al., PHENIX: building new software for automated crystallographic structure determination. Acta Crystallogr D Biol Crystallogr 58, 1948–1954 (2002).

71. L. Chi et al., The Dorsoventral Patterning of Human Forebrain Follows an Activation/Transformation Model. Cereb Cortex 27, 2941–2954 (2017).

72. D. Hockemeyer et al., Efficient targeting of expressed and silent genes in human ESCs and iPSCs using zinc-finger nucleases. Nat Biotechnol 27, 851–857 (2009).

73. A. E. Sullivan, S. D. M. Santos, An Optimized Protocol for ChIP-Seq from Human Embryonic Stem Cell Cultures. STAR Protoc 1, 100062 (2020).

74. P. J. Skene, S. Henikoff, A simple method for generating high-resolution maps of genome-wide protein binding. Elife 4, e09225 (2015).

75. J. D. Buenrostro, P. G. Giresi, L. C. Zaba, H. Y. Chang, W. J. Greenleaf, Transposition of native chromatin for fast and sensitive epigenomic profiling of open chromatin, DNA-binding proteins and nucleosome position. Nat Methods 10, 1213–1218 (2013).

76. J. D. Buenrostro et al., Single-cell chromatin accessibility reveals principles of regulatory variation. Nature 523, 486–490 (2015).

77. S. Chen, Y. Zhou, Y. Chen, J. Gu, fastp: an ultra-fast all-in-one FASTQ preprocessor. Bioinformatics 34, i884–i890 (2018).

78. B. Langmead, S. L. Salzberg, Fast gapped-read alignment with Bowtie 2. Nat Methods 9, 357–359 (2012).

79. H. Li et al., The Sequence Alignment/Map format and SAMtools. Bioinformatics 25, 2078–2079 (2009).

80. Y. Zhang et al., Model-based analysis of ChIP-Seq (MACS). Genome Biol 9, R137 (2008).

81. Q. H. Li, J. B. Brown, H. Y. Huang, P. J. Bickel, Measuring Reproducibility of High-Throughput Experiments. Ann Appl Stat 5, 1752–1779 (2011).

82. S. Heinz et al., Simple combinations of lineage-determining transcription factors prime cis-regulatory elements required for macrophage and B cell identities. Mol Cell 38, 576–589 (2010).

83. F. Ramírez et al., deepTools2: a next generation web server for deep-sequencing data analysis. Nucleic Acids Research 44, W160–W165 (2016).

84. D. Kim, J. M. Paggi, C. Park, C. Bennett, S. L. Salzberg, Graph-based genome alignment and genotyping with HISAT2 and HISAT-genotype. Nature Biotechnology 37, 907-+ (2019).

85. Y. Liao, G. K. Smyth, W. Shi, featureCounts: an efficient general purpose program for assigning sequence reads to genomic features. Bioinformatics 30, 923–930 (2014).

86. M. I. Love, W. Huber, S. Anders, Moderated estimation of fold change and dispersion for RNA-seq data with DESeq2. Genome Biology 15, (2014).

87. G. C. Yu, L. G. Wang, Y. Y. Han, Q. Y. He, clusterProfiler: an R Package for Comparing Biological Themes Among Gene Clusters. Omics 16, 284–287 (2012).

88. R. K. Smith-Bolton, M. I. Worley, H. Kanda, I. K. Hariharan, Regenerative growth in Drosophila imaginal discs is regulated by Wingless and Myc. Dev Cell 16, 797–809 (2009).

